# Going New Places: Successful Adaptation and Genomic Integrity of Grain Amaranth in India

**DOI:** 10.1101/2024.09.04.610786

**Authors:** Akanksha Singh, Markus G Stetter

**Author notes:** Dept. of Plant Sciences, University of Cologne, Cologne, Germany.

## Abstract

Global climate change will impact worldwide crop yields, requiring shifts and adaptation of crop varieties. The recent global spread of crops across different continents shows how plants successfully colonized new environments. One such spread is the introduction of the nutritious pseudocereal amaranth to India. Grain amaranth has been domesticated over 6,000 years ago in three different regions of the Americas and was introduced to India approximately 500 years ago. Nowadays numerous local landraces grow throughout the country’s wide climatic conditions. We investigate the introduction of grain amaranth to India to understand the factors allowing successful establishment of crops to novel environments, using whole genome sequencing of almost 200 accessions from India and more than 100 accessions from the crop’s native distribution. We find comparable levels of genetic diversity in the Americas and in India, despite the likely population bottleneck during the introduction to India. Surprisingly, the three grain amaranth species that were introduced do not show signs of gene-flow in India, while gene-flow in the Americas was high during the domestication of the crops. Correspondingly, the genetic differentiation between grain species was higher within India than in the native range, indicating a strong isolation between otherwise interbreeding populations. The reconstruction of the population history through demographic modelling of different scenarios suggested rapid expansion in the Indian population but a strong bottleneck in the native population, explaining the comparable diversity and isolation. We identified genomic loci under selection and associated with the climate in India that potentially enabled the adaptation to the new environment. These loci are predicted to provide an advantage under future climate scenarios, even in the native range. Our results suggest that introduced crops can act as reservoirs of genetic diversity, providing additional adaptive potential and resilience to future environmental change.

## Introduction

Rapidly changing environments pose major challenges for plant populations and crop production (Sloat et al. 2020). The adaptive potential of an organism is largely determined by its genetic compositions, i.e., standing genetic variation, new mutations and gene-flow (Excoffier et al. 2009; Exposito-Alonso et al. 2018; Waldvogel et al. 2020). Disentangling the relative contribution of each factor may help in better understanding of mechanisms of rapid adaptation.

The introduction of a species to a new region is a particularly strong change that often requires adaptation and can even lead to speciation (Irimia et al. 2021; Schluter 2001). Range expansion can lead to changes in population diversity and often requires rapid adaptation (Excoffier et al. 2009). Understanding the genetic consequences of range expansion is important to mitigate the impact of progressing rapid climate change, where most of the species are expected to migrate for new, potentially suitable areas (Waldvogel *et al*. 2020), including crops (Sloat *et al*. 2020). The recent spread of crops around the globe created locally adapted populations that have diverged from their relatives in the native range (Takou *et al*. 2024; Brandenburg *et al*. 2017; Bellucci *et al*. 2023; Gutaker *et al*. 2020). The movement of crops allows to study rapid range expansion and adaptation (Gutaker and Purugganan 2023). Understanding the successful migration and local adaptation of crops in the past might reveal templates to enable species conservation and crop improvement.

Before their spread around the globe, crop species have been affected by various evolutionary forces during their domestication (Purugganan 2019). Domestication has often been associated with increased genetic drift due to domestication bottlenecks (Wang *et al*. 2017; Stetter *et al*. 2020; Hyten *et al*. 2006; Caicedo *et al*. 2007). Selection for domestication traits also contributed to decreased genetic diversity in crops, compared to their wild relatives. Hence, crop populations potentially have less standing genetic variation left for future selection to act upon during range expansion (Beissinger *et al*. 2018; Stetter 2020). The spread of crops to new locations thus enables to study adaptation in populations that experienced recent directional selection and demographic changes. However, the spread and divergence of crops to distant locations outside their native range has only received little attention. (Bellucci et al. 2023; Brandenburg et al. 2017; Bančič et al. 2024; Takou et al. 2024). Little is known about functional diversity that enabled crops to establish and adapt to new environmental conditions in introduced locations. Reconstructing the patterns of their spread and the adaptive divergence could help to predict future capabilities of crops and natural populations.

Grain amaranth is a nutritious pseudo-cereal from the Americas that has been domesticated in the Americas and is now grown across the world, particularly in Africa and Asia. The crop could gain importance in the future due to its gluten-free nature, its high protein content, balanced micronutrient and rich antioxidant content (Joshi et al. 2018). Amaranth has been domesticated three times in different regions; twice in Mesoamerica (*A. hypochondriacus* L. and *A. cruentus* L.) and once in South America (*A. caudatus* L.) from one ancestral species (*A. hybridus* L.) (Stetter et al. 2020). While it was of high importance for the Aztecs and Incas, nowadays grain amaranth cultivation has strongly declined since the arrival of the Spanish in the 15th century (Brenner et al. 2000). The largest global exporter of Amaranth today is India (Santra et al. 2024; Brenner et al. 2000). Despite its recent arrival in India within the last 500 years, grain amaranth has become an integral part of cultural traditions and is widely grown across the country’s wide climatic conditions (Singh 2017; Brenner et al. 2000). The multiple domestication in different regions and the joint introduction into India provides the opportunity to understand the relative contribution of starting populations to the introduced crop and their involvement in its successful establishment 1).

In this study, we use whole genome sequencing of over 300 amaranth accessions, collected in the native range of grain amaranth and from India to identify genetic changes allowing successful establishment of crops to a novel environment. We find that the genetic diversity of introduced populations was comparable to those in the native range, despite an introduction bottleneck. Through demographic modeling, we attribute this to a strong population decline in the native range and rapid expansion in introduced populations after their split. Despite their gained geographic overlap of the three grain amaranth species we find very high genetic isolation between them and no signals of introgression. We identified several loci selected in India that are potentially involved in the local adaptation of these populations after their arrival to India. Overall, our results suggest that the introduction to India rescued genetic diversity that was lost in the native range that could provide adaptive potential and resilience to future environmental change even in the native range of the crop.

## Results

### Population structure is maintained in the introduced range recapitulating domestication history of amaranth

We sequenced 190 new amaranth accessions to an average coverage of 4.3x (min. 0.4x - max. 11.1x) and analysed whole genome sequences of more than 300 accessions, representing the three domesticated grain amaranth species and their wild relatives. Our sample includes diverse accessions of grain amaranth from the Americas (‘native range’) as well as the recently colonized India (‘introduced range’), allowing us to study the genetic signatures of recent plant expansion (Figure2A and Supplementary table S1). All samples had high mapping quality with over 98.43% of reads mapped to the *A. hypochondriacus* reference genome (Winkler et al. 2024). We identified a total of 13.63 million bialleleic SNPs after filtering for quality.

The repeated domestication of amaranth in different regions of the Americas led to geographic barriers between the crop species (Stetter *et al*. 2020), yet the three species showed signatures of gene-flow in their native range (Gonçalves-Dias *et al*. 2023). As all three species have been introduced into the same geographic areas of idea, strong intermixing between species might be expected (Figure 1A). The Principal Component Analysis (PCA) revealed three major clusters representing the three domesticated grain amaranth species as present in the native range. The Indian grain amaranth accessions clustered with the accessions of the respective species from the native range (Figure 2B). The ADMIXTURE analysis further confirmed the clustering pattern and absence of global patterns of admixture among the different species within India (Supplementary Figure S1). Despite the introduction of all three species into overlapping geographic regions of India, grain amaranths have maintained their global species identity.

**Figure 1.**
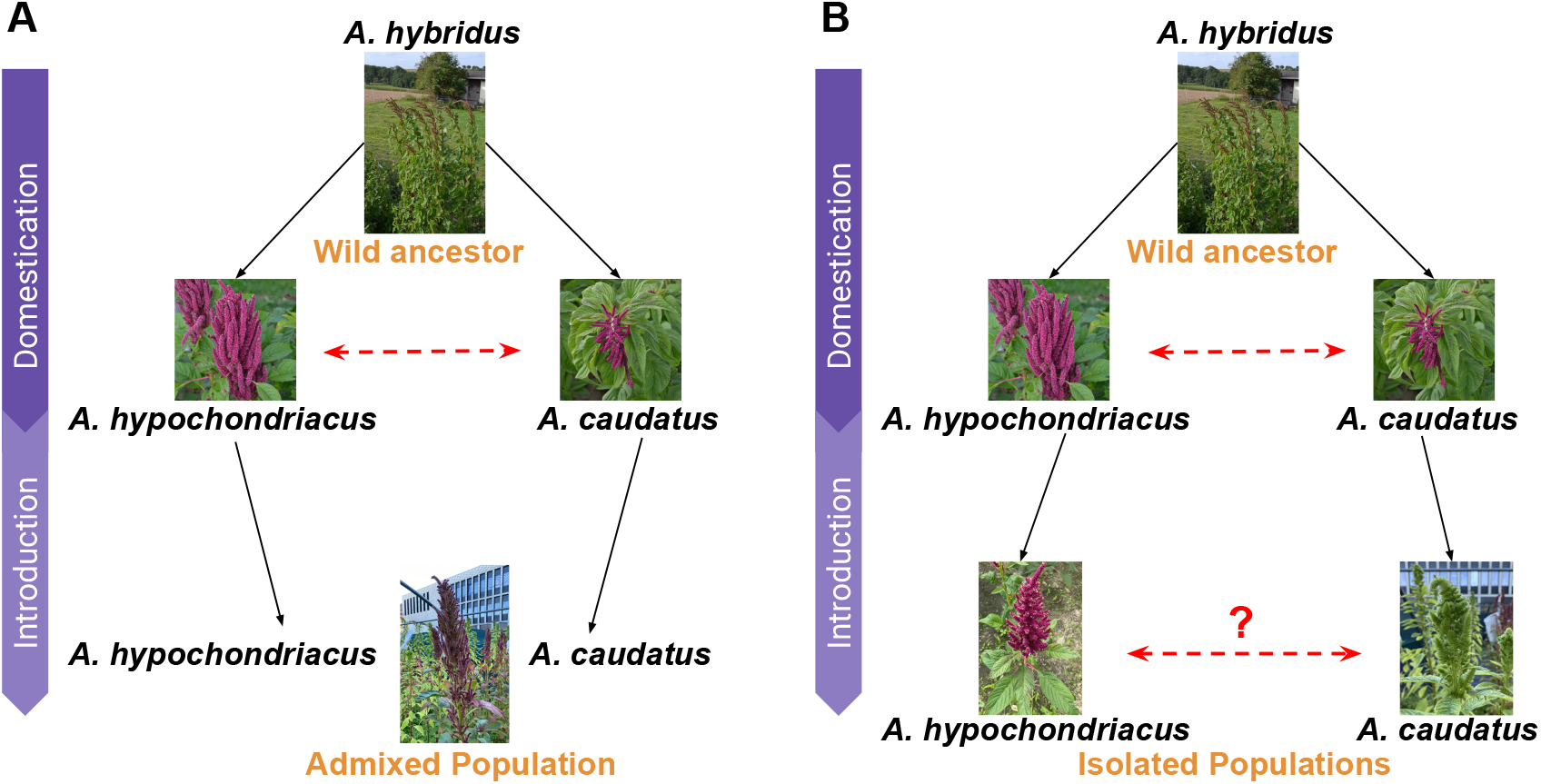
Schematic scenarios for crop introduction. Grain amaranth was domesticated in two regions (*A. hypochondriacus* in Central America and *A. caudatus* in South America) from the common wild ancestor (*A. hybridus*). Previous work has shown introgression between the two species in the native range ((Gonçalves-Dias et al. 2023) (depicted by red dotted arrow). The introduction to the new range (India) might have proceeded (A) through hybridization and admixture between the two crop species leading to a strongly admixed population; (B) the two species remained apart from each other showing high genetic differentiation with little or no gene-flow shown by red dotted line with a question mark.

**Figure 2.**
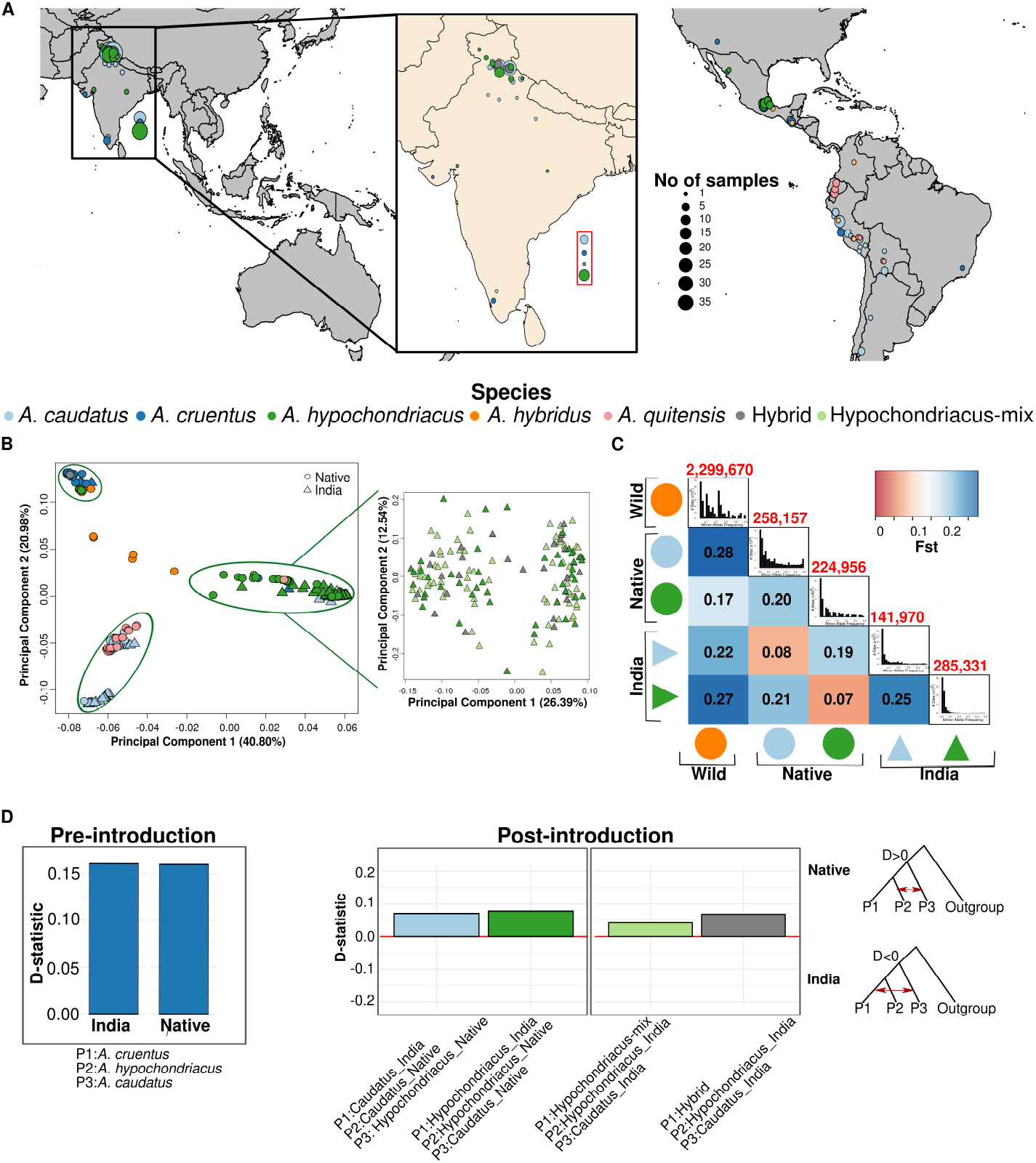
Population structure of Indian grain Amaranth. (A) Map of collection locations of accessions used in the study. Inset shows a zoom-in of the introduced range (India). The size of the dots represents the sample number per location. Samples represented in the red box in the Indian ocean had no associated geographic coordinates. (B) PCA of all individuals. Inset shows PCA with only *A. hypochondriacus*, Hypochondriacus-mix and Hybrid. (C) Heatmap representing F_ST_ statistic between each pair of populations. The diagonal shows 1D-site frequency spectrum (SFS) of each population with the numbers above representing populationspecific private alleles. (D) Gene-flow (ABBA-BABA) between grain amaranth populations. Pre-introduction represents gene-flow that occurred before the introduction to India, while post-introduction represents potential gene-flow after the introduction to India. The D-statistic indicates the strength of gene-flow. Positive D-statistic is gene-flow between P2 and P3 (representing gene-flow in the native region), while negative is between P1 and P2 (gene-flow in India).

Although the genetic distinction between groups was high, a number of accessions (74) taxonomically classified as *A. caudatus* and ‘hybrid’, strongly clustered with the *A. hypochondriacus* (renamed according to new cluster in Supplementary table S1). These accessions could not be distinguished from *A. hypochondriacus* even comparing higher PCs, as well as through a PCA of only these accessions with *A. hypochondriacus* (Figure 2B (inset), Supplementary Figure S2). Despite their high genetic similarity we called these populations ‘Hypochondriacus-mix’ and ‘Hybrid’, respectively, and retained them as separate populations in consecutive analyses because of the discordance between taxonomic classification and genetic identity. This discordance was surprising, given the strong species differentiation, but provided an unbiased sample for gene-flow analysis, as the groups are morphologically indistinguishable.

The strong species identity in India was confirmed by the mean fixation index (F_*ST*_) between the two grain species (*A. caudatus* and *A. hypochondriacus*) which was 0.25, 25% higher than in the native range (F_*ST*_ =0.20, Figure 2C). The mean differentiation between populations of the same species in the two regions was low (0.08 for *A. caudatus* and 0.07 for *A. hypochondriacus*). The greater differentiation between wild and introduced crop species compared to the differentiation between the wild and native crop populations provides additional evidence of introduction of grain amaranth in India from the native range after domestication. Further, mean F_*ST*_ between ‘Hypochondriacus-mix’ and *A. hypochondriacus* from India was very low (F_*ST*_ =0.006), while it was higher between ‘Hypochondriacus-mix’ and *A. caudatus* from India (F_*ST*_ =0.27), providing additional evidence for unambiguous genetic similarity of ‘Hypochondriacus-mix’ with *A. hypochondriacus* (Supplementary figure S3A).

Despite the high differentiation and the lack of population homogenization in India, smaller segments of the genome might have been exchanged between populations to improve adaptive potential (Figure 1B). Across the Americas gene-flow between amaranth species was prevalent and shaped modern grain amaranth populations (Gonçalves-Dias *et al*. 2023). Contrary to the strong gene-flow in the Americas, we observed no genome-wide signals of gene-flow within the Indian grain amaranth species, despite the geographical proximity of accessions in India (Figure 2D). We further estimated local ancestry for each individual from India using native species group as ancestors employing Efficient Local Ancestry Inference (ELAI) (Zhou *et al*. 2016). Our results were in close agreement with the previous results of no admixture among the groups, where > 99% of the genome was from the respective species (Supplementary Figure S4).

We also could not identify any prevalent admixed loci in the Hypochondriacus-mix and Hybrids, both of which were purely assigned to *A. hypochondriacus* genome unambiguously (Supplementary Figure S4). Given the obvious difficulty to taxonomically distinguish *A. caudatus* and *A. hypochondriacus* but the lack of genetic differentiation, the species determining characteristics might be controlled by a limited number of loci. Hence we used the previously classified taxonomic species as phenotype for a genome wide association study (GWAS) and identified 22 significant associations (Supplementary figure S5). Within 10kb upstream and downstream of the significant SNPs were annotated with 14 genes (Supplementary table S2), that could control for different developmental features in plants and might be involved in flower morphology which is being used to delimitate the species.

### High genetic diversity in Indian amaranth

The introduction of populations to new locations often leads to a reduction in genetic diversity. Such a reduction might have severe consequences for crops, as they experienced a recent reduction in diversity during their domestication. The strong reduction of diversity during a new colonisation might be mitigated by gene-flow, which is very limited in Indian grain amaranth. To understand the consequences of the introduction for genetic diversity, we calculated the number of private alleles per population and compared them between populations (Figure 2C). As expected the highest number of private alleles was observed in the wild ancestor *A. hybridus*. However, the number of private alleles of both grain amaranth species *A. hypochondriacus* and *A. caudatus* were similar in India and the native range (Figure 3A-B). The wild *A. hybridus* also had the highest nucleotide diversity (= 8.5E-3), followed by the three domesticated species. However, we did not observed further reduction of the genetic diversity in the crop species from India as compared to those from the native range (Figure 3C and Supplementary table S3). Further, the Tajima’s D value were negative for all the domesticated grain amaranth species (Figure 3D). The lower negative Tajima’s D value for the introduced Indian range could be a potential indication of recent population expansion. The site-frequency spectra also indicated an excess of rare alleles in the introduced populations, together suggesting recent population growth after introduction (Figure 2C and Supplementary figure S3).

**Figure 3.**
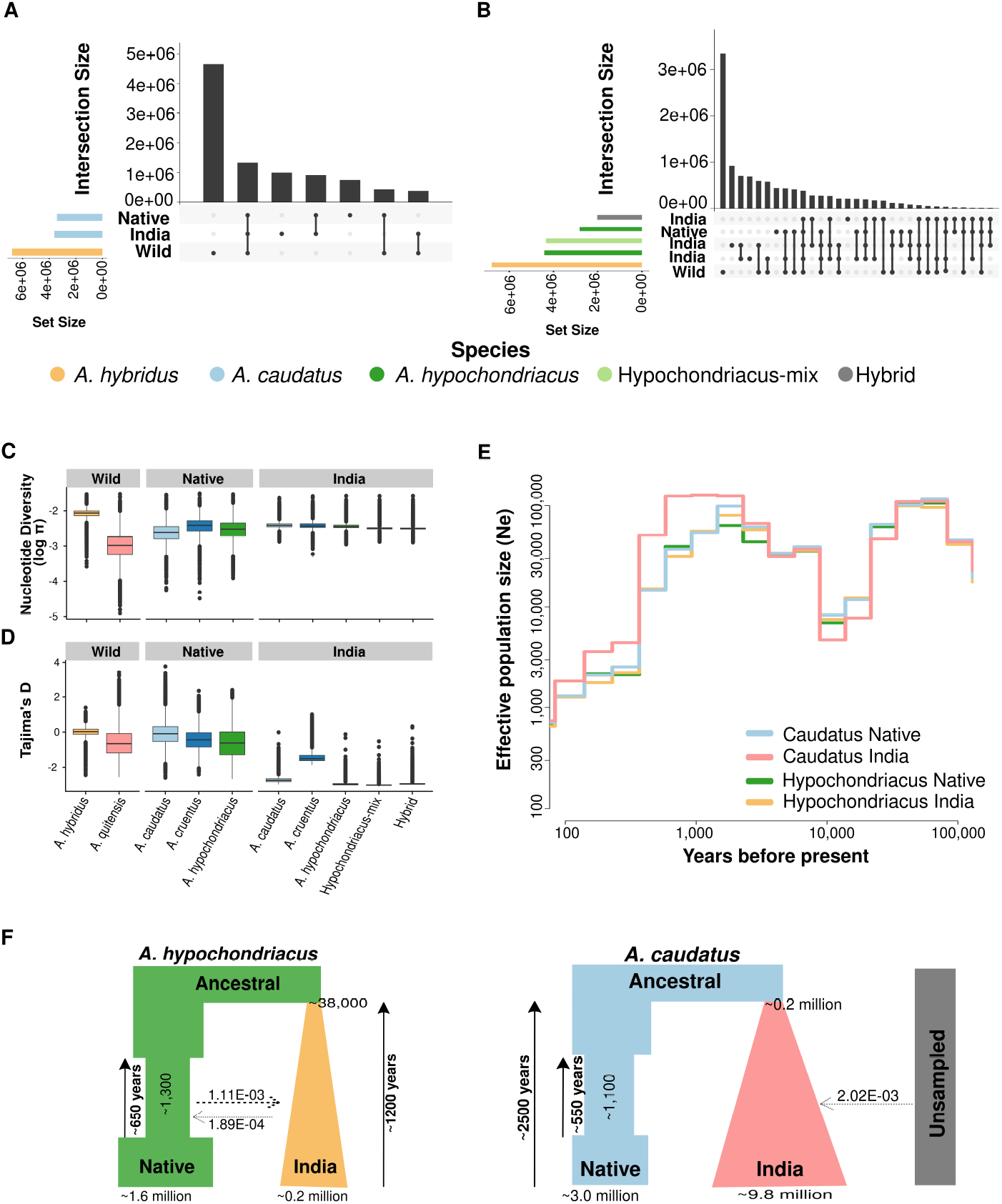
Genetic diversity and demographic history. (A -B) Number of private and shared alleles between two regions and native wild species for (B) *A. caudatus* and (B) *A. hypochondriacus*. (C-D) Distribution of (C) nucleotide diversity (log_10_ transformed) and (D) Tajima’s D in 10kb windows. (E) Population size estimate of the two species in native and introduced region showing bottleneck and expansion. (F) Best-fit demographic history model of introduction to India for the two species.

### Demographic history of introduction of grain amaranth to India

We aimed to reconstruct the history of introduction and model the population dynamics during the colonisation of India. We used popsizeABC to first predict the dynamics of effective population size for these population. All the populations indicated a domestication bottleneck, consistent with previous work (Stetter *et al*. 2020) (Figure 3E and Supplementary figure S6). We further employed simulations using Fastsimcoal2 (Excoffier *et al*. 2021) to model the introduction history with six alternative models with varying population sizes and different extend of migrations (Supplementary figure S7). We identified two different models that fit best for the two grain amaranth species (Figure 3F, Supplementary table S4). For *A. hypochondriacus*, a bottleneck in the native region after the split of the Indian population and population expansion in India along with the continuous geneflow from the native range might have led to the observed data.

Migration rates were asymmetric, with higher migration from the native range to India compared to the opposite. Similarly, the best fitting model for *A. caudatus* showed a bottleneck in native population with expansion in the Indian population and migration from an unknown population without any further contact with the native region after split. The estimated split time between the two populations was recent, aligning with a recent introduction of grain amaranth in India, even though the absolute number reaches back to over 1,000 years. However, estimating exact split times within a few hundred year range in such a complex model is difficult. The result shows that the introduction likely occurred recently. Together, these results show a recent introduction of grain amaranth to India that enabled the maintenance of high diversity in India, but also indicates the reduction of diversity in the native range.

### Region specific selective sweeps differ between species

To identify genomic regions under selection during the introduction of the crop, we performed a cross-population composite-likelihood ratio (XP-CLR) analysis between the native and introduced populations of both grain amaranth species (Chen *et al*. 2010). This enables the separation of selection before the introduction from potential local adaptation signals. Considering the top 1% genome-wide XP-CLR statistics as significance cut-off we identified 149 regions in *A. caudatus* and 146 regions in *A. hypochondriacus* under selection (Figure 4A). Only eight sweep regions were common between *A. caudatus* and *A. hypochondriacus* (p-value = 1.494852e-05). The Hybrid and Hypochondriacus-mix populations, showed strong overlaps with *A. hypochondriacus* enforcing the shared history of these populations (Figure 4B and supplementary figure S8).

**Figure 4.**
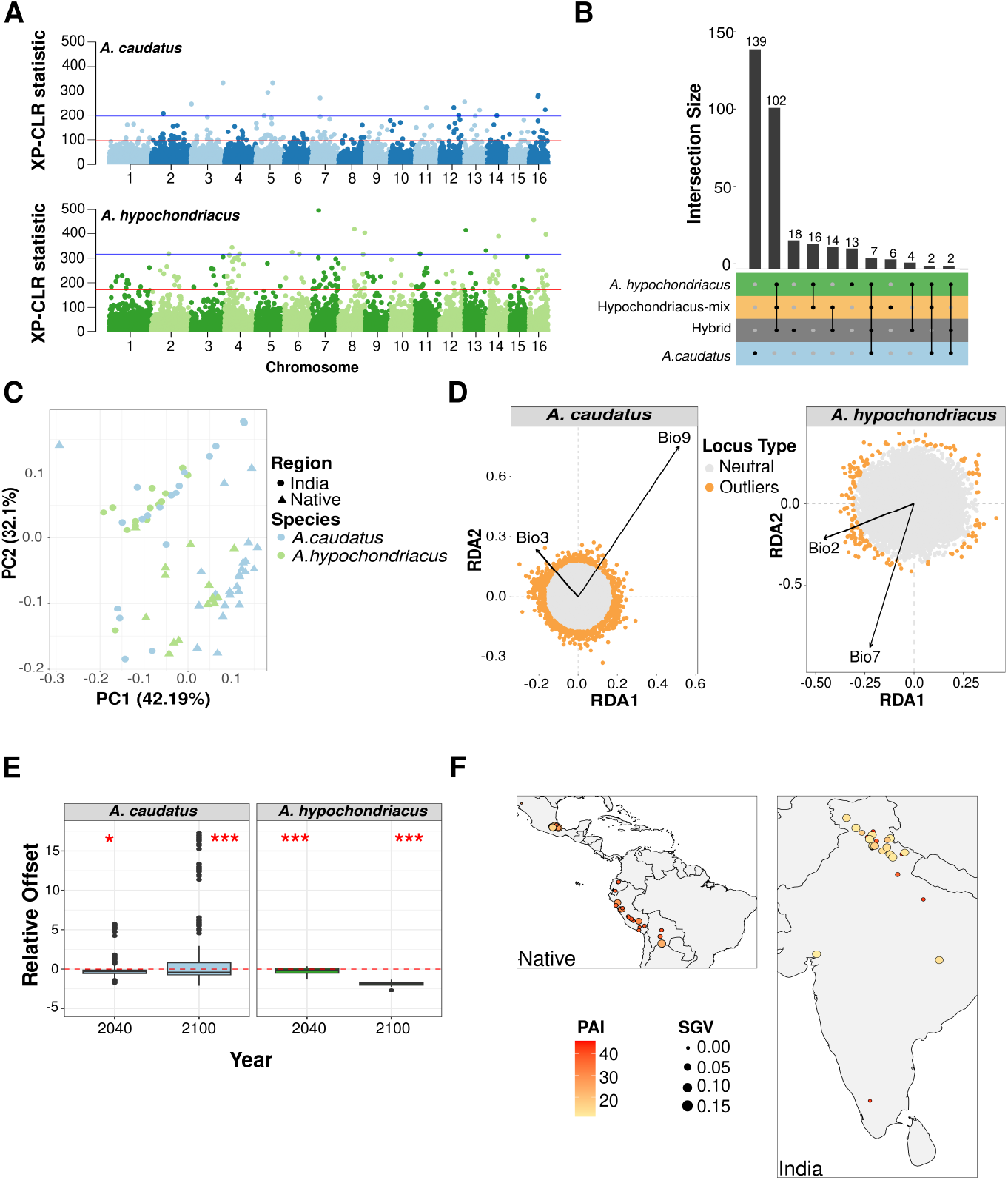
Putative selective sweeps, climatic associations and genomic offset. (A) Genome-wide distribution of XP-CLR statistic for the two species using their native counter-part as reference population. Red line represents top 1 percentile and blue line represents 0.1 percentile. (B) Shared putative selective sweep regions identified as top 1 percentile genome-wide XP-CLR outlier. (C) PCA conducted using 19 bioclimatic variables extracted from WorldClim database. (D) RDA plot representing association of candidate adaptive loci with the two significantly associated bioclimatic variables for the two species (Bio2 = Mean diurnal temperature range, Bio7 = Temperature annual range, Bio3 = Isothermality and Bio9 = Mean temperature of the driest quarter). Outliers were identified as loci with FDR <0.05. (E) Relative offset (genetic offset - geographic offset) calculated, using candidate adaptive loci. For genetic offset two future climatic scenario (2040 and 2100) were considered. Geographic offset was calculated as difference in adaptive index for samples from India placed in the native climatic conditions. Red dotted lines show where genetic offset is equal to geographic offset. Deviation towards positive and negative values represent lower and higher geographic offset, compared to predicted genetic offset, respectively. The asterisks above the box-plot represent significance level for one-sample t-test (* P-value<0.05, *** P-value<0.001). (F) Map showing the levels of Population Adaptive Index (PAI) and adaptive standing genetic variation in the sampled populations of the two species.

To find potential functions of selected regions we performed a GO analysis. The sweep regions in *A. caudatus*, overlapped with the 77 genes, enriched for three biological processes; cellular macro-molecule metabolic process, metabolic process and nucleic acid phosphodiester bond hydrolysis (Supplementary table S5). In *A. hypochondriacus*, 58 genes were annotated within sweep regions. These were enriched for the three GO terms; cellular macromolecule metabolic process, protein phosphorylation and peptide transport. Only three genes were found to overlap between the two species that were identified to be Omega-hydroxypalmitate O-feruloyl Transferase (AHp006427), NB-LRR family (AHp006793) and Regulator of nonsense transcripts 1 homolog (AHp011497). Sixteen and 26 of the XP-CLR candidate regions in *A. caudatus* and *A. hypochondriacus*, respectively were also identified as selective sweeps in Indian amaranth using RAISD (Supplementary figure S9). The selection analysis indicates that local adaptation in the different grain amaranth species proceeded through different genetic changes with only few major adaptive genes and potentially through soft selective sweeps and polygenic changes (Pritchard and Di Rienzo 2010).

### Local climate partly governs the observed genetic variations

Local climatic conditions might be a driver for observed genetic variability, leading to a correlation between climate and diversity. A PCA on 19 bioclimatic variables showed that the native region and India differ in their over all climatic conditions (Figure 4C). We further conducted Redundancy Analysis (RDA) to identify the climatic and spatial factors that structure the distribution of genetic variation and partition the variance explainable by the associated climatic variable. Using a forward selection model, we identified two climatic variables, mean diurnal range and temperature annual range (bio2 and bio7) explaining 13.6% of the total explainable genetic variation in *A. hypochondriacus*, while isothermality and mean temperature for driest quarter (bio3 and bio9) were explaining 10.8% of explainable variation in *A. caudatus* (Supplementary table S6 and S7). Using a multivariate constrained ordination technique with population structure correction of the RDA, we identified 178 and 729 loci significantly associated (FDR <0.05) with environmental variables in *A. hypochondriacus* and *A. caudatus*, respectively. We found 154 and 679 genes in *A. hypochondriacus* and *A. caudatus*, respectively within 20kb around associated loci, of which only two genes overlapped between the two sets (Supplementary table S8). However, five genes that were in climate associated regions overlapped with selective sweeps. This indicates that local climate might have driven selection in introduced grain amaranth.

### Populations from introduced ranges can act as potential restorer of native population diversity

We estimated that the current landscape adaptation explained approximately 43% of adaptive genetic variation in the two species (Figure 4D). To predict the potential of these genetic resources for future climatic adaptation, we estimate the genomic offset for two scenarios of early (2040) and late future (2100) climatic conditions (Supplementary figure S10). The observed genomic offset for native and the introduced range was not significantly different for *A. caudatus* for year 2040 (pvalue= 0.59), but was marginally higher in the native range for the year 2100 (mean genomic offset, India = 0.218, Native = 2.175, pvalue =0.057). However, for *A. hypochondriacus*, the offset was significantly higher in India than in the native range for both future scenarios of 2040 (pvalue=1.9e-05, native=0.45, India = 0.7) and 2100 (pvalue = 0.0004, native=1.06, India = 1.26). We further estimated the relative offset as difference between predicted genetic and geographic offset, where the values larger than zero represent a reduction in genetic offset in the native range when introducing populations from India. The relative offset was less than zero for *A. hypochondriacus* in both the future scenarios, but was significantly higher than zero for *A. caudatus* in the future scenario of 2100 (Figure 4E). Further, the populations in the introduced range showed higher standing genetic variation and a lower population adaptive index for the majority of populations (Figure 4F). These results together provide evidence that genetic variation from the introduced range could potentially provide adaptive advantages under future climates even in the native range.

## Discussion

We studied the introduction of grain amaranth to India as model for crop expansion in the last centuries. Our results underline the complexity of such spreads and the high potential of introduced crop populations as pre-adapted resources for future crop improvement.

The repeated domestication and strong signals of gene-flow between grain amaranths even over long distances in the Americas (Stetter *et al*. 2020; Gonçalves-Dias *et al*. 2023), provided high potential for a complete admixture between grain amaranth species in India where they are grown in overlapping geographic regions (Figure 1A). Yet, the analysis of population structure showed that the three grain amaranth species maintain high species identity, demonstrated by the grouping in the PCA, the high F_*ST*_ between populations in India and the lack of overlapping signals of positive selection (Figures 1B, 2, 4). A potential hypothesis for the maintained differentiation might be the intentional separation by humans even after introduction, however, the taxonomically ‘misidentified’ individuals of ‘Hypochondriacus-mix’ shows that human separation is complicated. These samples were taxonomically classified as *A. caudatus*, but genetically unambiguously cluster with *A. hypochondriacus*, without signals of gene-flow between them (Figure 2). These samples represent an ideal control, as intentional separation from *A. hypochondraicus*, but not from *A. caudatus* would have altered their genetic composition. Nonetheless, they show almost 100% *A. hypochondriacus* identity and no signs of *A. caudatus* introgression (Figure 2 and Supplementary figure S4). Another reason for the high differentiation might have been a lack of cross breeding, but outcrossing rates are up to 20 % (Stetter *et al*. 2016) and the high flower number and small seeds make selection against hybrids inefficient, which led to strong outcrossing between crops and even with wild relatives has been frequent in the Americas during the domestication of amaranth (Stetter *et al*. 2020). Previous work indicated that reproductive barriers evolved after domestication in the Americas through genetic incompatibilities between *A. cruentus* and *A. caudatus* (Gonçalves-Dias *et al*. 2023). Such barriers might have further evolved in India, preventing extended gene-flow between species. Given the incomplete speciation between *A. hypochondriacus* and *A. caudatus* in allopatry in the Americas, with ample gene-flow and fit offspring (Gonçalves-Dias *et al*. 2023), the strong separation between the two species in India might represent a case of secondary reinforcement (Dobzhansky 1982; Hopkins 2013). The partial isolation in allopatry between the two species likely progressed in sympatry in India, where the two species seem to not interbreed anymore (Figure 2). Further evidence on hybrid fitness in the two ranges would be required to understand the driver of reproductive isolation, which has rarely been identified in crops (Hopkins 2013). The strong reproductive isolation in India further indicates the ongoing speciation among the three grain amaranths, despite their high morphological similarity (Sauer 1967). The evolution of reproductive barriers in the relatively short time of domestication might be more frequent across crops species (Tenaillon *et al*. 2023).

The mixing of crop species in the new range would have led to an homogenization of allele frequencies across populations in India and an apparent increase in genetic diversity. The absence of admixture in India and the introduction-related bottlenecks could have decreased genetic diversity (Figure 3). Despite the strong population structure and demographic history, amaranth showed equally high genetic diversity in India as in the Americas (Figure 3). This is likely due to a continuous introduction of additional germplasm from the native range and other derived regions and a loss of genetic diversity in the native range (Brenner et al. 2000) (Figure 3). Similar observations have been made in common bean, where to Europe introduced germplasm harbours higher diversity than populations from the primary domestication center in the Andes (Bellucci et al. 2023). For amaranth such a decline might have partially resulted from the prohibition of amaranth cultivation, by the Spanish (Sauer 1967; Brenner et al. 2000), by a massive decline in native American population (O’Fallon and Fehren-Schmitz 2011) and the success of other crops in the Americas in the last centuries (Hancock 2022). Such strong changes in population sizes in the primary centers of a crop’s diversity, show the importance of introduced ranges for the conservation of genetic diversity as native ranges might loose further diversity due to rapidly changing environments. Our results show that potentially adaptive diversity differs between populations as the two different amaranth species that were introduced to India show distinct signals of selection despite their introduction to the same range (Figure 4B). Hence, they might provide different pre-adaptations for future climatic conditions and be a well suited resource for plant breeding. In addition to crop wild relatives that are mostly found in the native center of domestication introduced crop populations are well adapted to diverse environments (Hao et al. 2020; Yang et al. 2023). Together they harbour additional variation that has potential to be utilised for crop improvements with a wide range of pre-adapted alleles for novel selection pressures (Muleta *et al*. 2022; Bhullar et al. 2010; Haupt and Schmid 2020).

Evolutionary convergence by human selection provide a channel of constraints that shape adaptation of crops for human use (Purugganan 2019). The presence of three independent domestications in grain amaranth showed parallel evolution of domestication-related traits with independent signals of selection (Stetter et al. 2020). The introduction to a similar environment in India also led to mostly independent signals of selection, but few of the genomic regions under selection were common to both species. This overlap was larger than expected by chance, suggesting that few major genes might show a similar response to local selection pressure in the introduced range. Similar results have been found in other crops (Zhao et al. 2023; Bellucci et al. 2023; Takou et al. 2024). The overlapping genes in response to the introduction to India are involved in regulating biotic and abiotic stress in *A. thaliana* (Gou et al. 2009; Shi et al. 2012), suggesting the requirement of mitigating stressful environments after the introduction. Yet, such major genes remain the exception, reinforcing the growing evidence that local adaptation is highly polygenic (Buckler et al. 2009; Tyrmi et al. 2020; Haupt and Schmid 2022). We also identified significant climatic association of several loci (Figure 4D), suggesting potential polygenic and convergent environmental adaptation, also found in other species (Zhao et al. 2023; Luqman et al. 2023; Haupt and Schmid 2022). In the presence of standing genetic variation, polygenic adaptation can proceed rapidly, as only small allele frequency changes are required to achieve large trait changes (Jain and Stephan 2017; Stetter *et al*. 2018). Given the polygenic nature of climate adaptation, maintenance of high variability of the adaptive loci is important to maintain and increase future population adaptability, supporting assisted migration (Broadhurst *et al*. 2008; Hung *et al*. 2023). Locally adapted populations from around the globe might provide such standing genetic variation that can enable the maintenance of high crop performance.

Together, our results show the complexity of rapid plant expansion to distant environments. The successful introduction of domesticated plants beyond their native range is mediated by several genetic factors and can proceed without gene-flow. Selection and population histories might even lead to speciation and genetic incompatibilities. Overall, introduced populations can harbour high genetic diversity and alleles that have been lost in native populations. These alleles might be able to mitigate the negative effects of changing environments on crop production and can serve as reservoir of pre-adaptation.

## Materials and Methods

### Plant material and sequencing

We accessed all available seeds of grain amaranth species tagged with country of origin as “India” from USDA-ARS genebank. These 190 accession included all three grain amaranth species, *A. caudatus* L. (106), *A. cruentus* L. (5) and *A. hypochondriacus* L. (53) and few accessions marked as “hybrids” (26) due to taxonomic veracity (Supplementary table S1). We grew plants in the greenhouse and extracted DNA from young leaf tissue using NucleoSpin Plant II kit (Macherey-Nagel) according to manufacturer’s instructions. We used modified Nextera-shallow sequencing protocol described in Rowan *et al*. (2019) to make the libraries. Briefly, we mixed 2uL of DNA (conc. 0.5ng/uL) with 2.1 uL of sterile water, 0.818uL tagmentation buffer and 0.0818uL tagmentation enzyme from Illumina Tagment DNA Enzyme and Buffer Kit (Cat. No. 20034197). We incubated the samples at 55^°°B°;^C for 10min and then allowed to cool at room temperature. Then we added 5uL of custom index and barcoding primers and used KAPPA2G Robust PCR kit with GC buffer (Cat. No. KK5004) for PCR amplification. We amplified the samples using PCR amplification cycle as: 72^°^C for 3 min, 95^°^C for 1 min, 14 cycles of 95^°^C for 10sec, 65^°^C for 20sec, and final extension at 72^°^C for 3min and then checked the library success and size distribution on 2% agarose gel. We further pooled 5uL from each of the successful libraries followed by dual size selection (300-500bp) using Promega Pronex-Size Selective Purification System (Cat.No. NG2001) following manufacturer’s instructions. The pooled libraries were sequenced on Illumina NexSeq with 2×150 bp by Novogene, United Kingdom. In addition to Indian grain amaranth accessions, we also downloaded raw fastq files for 120 accessions (34 of *A. caudatus* L., 25 of *A. cruentus* L., 24 of *A. hypochondriacus* L., 9 of *A. hybridus* and 25 of *A. quitensis* from the native range from European Nucleotide Archive (project number: PRJEB30531) (Stetter *et al*. 2020). However, for the final in-depth analysis, we used only 88 unambiguously taxonomically and genetically classified accessions from the native range in addition to the new Indian accessions (Gonçalves-Dias and Stetter 2021). *A. tuberculatus* was used as outgroup (ERR3220318) (Kreiner *et al*. 2019).

### Mapping and variant calling

We mapped both sequenced and downloaded reads to the *A. hypochondriacus* reference genome V2.2 (Winkler et al. 2024) using BWA-mem2 (V2.2.1) (Vasimuddin et al. 2019). We further downsampled the downloaded samples with very high sequencing depth to match those sequenced in the study. We used Samtools (V1.13) (Li et al. 2009) to sort bam files and Picard tools (https://github.com/broadinstitute/picard) to mark the duplicates. We used ANGSD (v.0921) (Korneliussen et al. 2014) to call variants using flags -gl 2 -dovcf 1 and applying quality filters as -remove-bads 1, -minMapQ 30, -only_proper_pairs, -trim 0, and SNP_pval 1e-6. After assessing the quality, we removed individuals with more than 80% sites as missing (13 individuals) and re-called the variants from a total of 268 individuals (Supplementary table S1). For further filtering we used VCFTOOLS (v.0.1.17) (Danecek et al. 2011) using -max-missing 0.8, -minDP 3 and -maf 0.002, resulting into 13 million SNPs. We used PLINK (Purcell *et al*. 2007) for LD pruning which resulted into approx. 4 million LD-pruned biallleic SNPs.

### Population genetic analysis

We used Angsd (v.0921) to calculate population genetic summary statistics, i.e., site frequency spectrum (SFS), observed heterozygosity, nucleotide diversity and Tajima’s D. We also estimated 95% confidence interval of the nucleotide diversity of each population by calculating stats 50 times using Angsd by randomly sub-sampling 10 individuals from each population. LD filtered data was used to conduct Principal component analysis (PCA) and population structure analysis using PLINK and ADMIXTURE (Alexander et al. 2009), respectively. We re-assigned the accessions into their new genetic cluster based on the results of PCA (Supplementary table S1). We used VCFTOOLS to estimate the pairwise differentiation (F_ST_) between each genetic clusters from native and introduced range. To calculate number of private alleles per population (cluster) we used SnpSIFT (v5.1) private (Ruden et al. 2012) and used PopLDdecay (v3.42) (Zhang et al. 2019) to estimate linkage disequilibrium.

### Signals of gene-flow and local ancestry

We inferred genome-wide gene-flow between populations using D-statistic (ABBA-BABA) implemented in Angsd (v.0921) using abbababa2. *A. tuberculatus* was considered as outgroup. Further, we also predicted ancestry for every nucleotide position in each individual from introduced populations using local ancestry inference method ELAI (Zhou et al. 2016). For this we first identified un-admixed individuals from the native range using ADMIXTURE at predefined K=2 and K=3, corresponding to three grain amaranth species. Individual with ancestry probabilities in their own cluster of >0.99 were constituted as the source population species group. We ran ELAI using three upper-level and fifteen lower-level clusters with 30 EM steps. The time of mixing was set to 100 generations with filter parameters: “–exclude-nopos –exclude-miss1 –exclude-maf 0.01”. We analysed each chromosome separately with 10 replicated runs. Dosage of ancestry for each site was calculated as the mean of all the replicated runs. For identifying the loci associated with the taxonomic classification of the species we used CMLM model in GAPIT (Zhang et al. 2010). We used first three PCs and kinship matrix as control for population structure. We included accessions from *A. caudatus* and *A. hypochondriacus* from the native and introduced range using the previously classified taxonomic species as phenotype.

### Demographic analysis

We inferred historical effective population size of each population using PopSizeABC (Boitard *et al*. 2016) which uses local linkage pattern and alleles frequency to predict demography for recent times. To get detailed estimate of the most suitable scenario for the demographic history of introduction and establishment of grain amaranth into India, we used simulations in Fastsimcoal2 (Excoffier *et al*. 2021). We used a python program easySFS (https://github.com/isaacovercast/easySFS) to estimate the joint site frequency spectrum (SFS) using non-coding SNPs. First, we used the preview mode (–preview) to identify the true sample size and then projecting (-proj) the best sample size to generate the joint SFS. We tested six different base models namely, (i) simple two-population split, (ii) two-population split with expansion in Indian population, (iii) two population split with bottleneck in the native population, (iv) two population-split with bottleneck in native and expansion in Indian population,(v) two population split with decline in native population, and (vi) two population split with decline in native population and expansion in Indian population (Supplementary figure S7A). To these six base models we then added different migration scenarios; continuous migration, one time migration, two-time migration and finally migration from an unknown un-sampled population (Supplementary figure S7B). This unknown population was included in the model to comment on any unknown introgression from potential local pre-adapted wild population providing beneficial variation aiding in establishment of these populations in the introduced range. We ran 100 independent runs with 200,000 coalescent simulations and 40 cycles of likelihood maximization algorithm to estimate best parameters for each scenario. We identified best model as the one having the lowest Akaike’s Information Criterion (AIC). To estimate 95% confidence interval of the parameters for the best fitted model we used 50 non-parametric bootstrapping datasets. Each of the 50 bootstrapped datasets was run 100 times to estimate the best run. We used these 100 best runs to estimate the confidence interval using boot package in R (Canty *et al*. 2017).

### Selective sweep in introduced populations

We identified putative selective sweeps specifically in the introduced population after introduction to India using XP-CLR (Chen *et al*. 2010). We compared each population in India to its native population pair individually, which allowed us to identify putative candidate selective sweeps that are specifically under selection in the introduced range nullifying the effect of any selection already happened in the native range. We estimated the XP-CLR statistic for each chromosome in a window-size of 10kb, limiting to the maximum number of SNPs as 500 and ld of 0.8. We considered top 1% of the genome-wide XP-CLR statistic for each population pair as significant. In addition we also identified population specific selective sweep using default parameters in RAiSD (Alachiotis and Pavlidis 2018).

### Climatic differentiation, redundancy analysis and genomic offset

We used landscape genomics to identify the influence of local climatic conditions on the observed genetic variation. The association of the climate with the genetic variation could provide a clue for local adaption in the populations, provided the climate is significantly variable. Therefore, we first used PCA to visualize the overall climatic differentiation between the two regions (native and introduced). For these analyses, we only used the accessions for which the geographical coordinates or location data was available. We downloaded 19 bioclimatic variables for the near current data from WorldClim database (https://www.worldclim.org/data/index.html) at highest available resolution of 30 seconds which corresponds to 1 km^2^. Raster R package (Hijmans et al. 2016) was used to extract climatic data for each site using geographic coordinates.

We used Redundancy Analysis (RDA), to explore if and how spatial and environmental factors structure the amount and distribution of genetic variation among these populations (Forester et al. 2018). RDA is a multivariate constrained ordination method capable of analysing genotype-environment associations performing better than univariate approaches and random forest making it well suited to identify the loci under weak polygenic selection (Forester et al. 2018). We performed RDA individually for the two species (*A. caudatus* and *A. hypochondriacus*), following the guidelines described in Capblancq and Forester (2021). To exclude false association, we used LD pruned biallelic SNPs having minor allele frequency >5%. As some samples share same location and coordinates, we grouped them into sub-populations and calculated allele frequency for each sub-population and each genomic location. Since RDA does not allow any missing data we filtered out sites with more than 10%of missing data and imputed the rest with the median across the complete sampling. We used standardized current climatic data for 19 bioclimatic variables as predictor variable and first three principal components of PCA conducted using intergenic biallelic SNPs as proxy for correcting for the population structure. We used forward selection model (ordiR2step) with 1000 permutations in Vegan R package (Oksanen et al. 2022) to identify environmental variables explaining the significant amount of explainable genetic variance. Correspondingly, several partial redundancy analyses were then conducted to partition explainable genetic variance into geographical (latitude, longitude and elevation), neutral genetic variation and environment components. We also identified loci underpinning the local environmental adaptation using environmental variables showing significant explainable variance conditioned on population structure.

To maintain stringency and reduce false positives, we ran a similar genome-environmental scan without using population structure and overlapped them with the previously identified loci and considered only the overlapping loci for further prediction of adaptive landscape and genomic offset (Supplementary figure S11). We predicted genomic offset for future climate as euclidean distance between the present and future optimal adaptive genetic composition, considering the adaptive landscape for the present genotype with environment association. It gives an estimate of the amount of genetic change required to tract the future climatic conditions. We used two time frames in future climatic scenario for 2021-2040 and 2080-2100 for the prediction criterion of maximum carbon emission ssp585 under MRI-ESM2-0 general circulation model (https://www.worldclim.org/data/cmip6/cmip6_clim30s.html). To further predict the genetic performance of the populations in the home-vs-away concept, we also calculated spatial (geographical) genomic offset between the climate of source population and climate of the projected area for both present and future climates. Finally, we estimated the levels of standing genetic variation (SGV) in the populations and Population Adaptive Index (PAI) as described in Capblancq *et al*. (2020). Briefly, SVG was calculated as mean of allele frequency within population for the outlier adaptive loci and PAI was estimated as absolute allele frequency difference between population-specific and mean allele frequencies through all the populations. We used these parameters as estimate of adaptive capacity of populations, where SVG represents availability of adaptive alleles and PAI represents the extremeness of genetic adaptation. Lower PAI and higher SGV represents higher adaptive capacity of the population (Capblancq *et al*. 2020).

## Data Availability

The raw data of samples sequenced in the study can be accessed from European Nucleotide Archive (ENA) under bioproject PR-JEB76492. All scripts used in the analysis are available at https://github.com/cropevolution/Introduction-of-Grain-Amaranth-to-India

## Acknowledgments

We thank David M. Brenner from Iowa State University and US National Plant Germplasm System (USDA-ARS) for the seeds used in the study. We also thank Roswitha Lentz for helping in sampling and all the lab members for fruitful discussion helping improve the manuscript. We acknowledge funding by the Deutsche Forschungsgemeinschaft (DFG, German Research Foundation) under Germany’s Excellence Strategy – EXC-2048/1 – Project ID 390686111 and grant STE 2654/4 to MGS by the DFG. We furthermore thank the Regional Computing Center of the University of Cologne (RRZK) for providing computing time on the DFG-funded (Funding number:INST 216/512/1FUGG) High Performance Computing (HPC) system CHEOPS.

## Author contribution

AS and MGS planned and designed the study. AS performed experiments and analyzed the data. AS and MGS wrote the manuscript. All authors read and approved the manuscript.

