## Supplementary_Information for "Going New Places: Successful Adaptation and Genomic Integrity of Grain Amaranth in India"

**Supplement**

**Supplementary Figures**

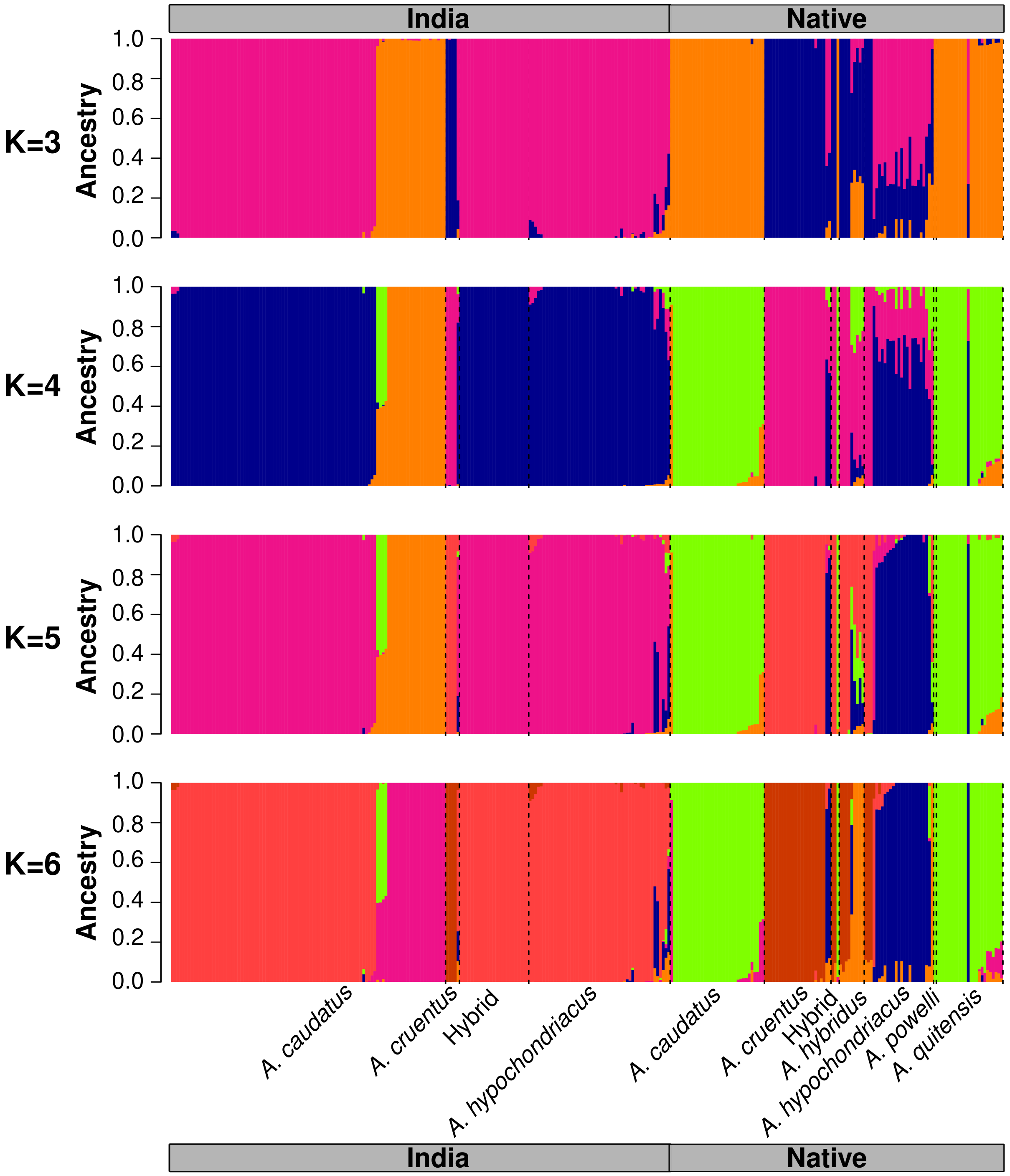


**Figure S1** Bar-plot representation of individual ancestry proportion estimated with ADMIXTURE for different K.

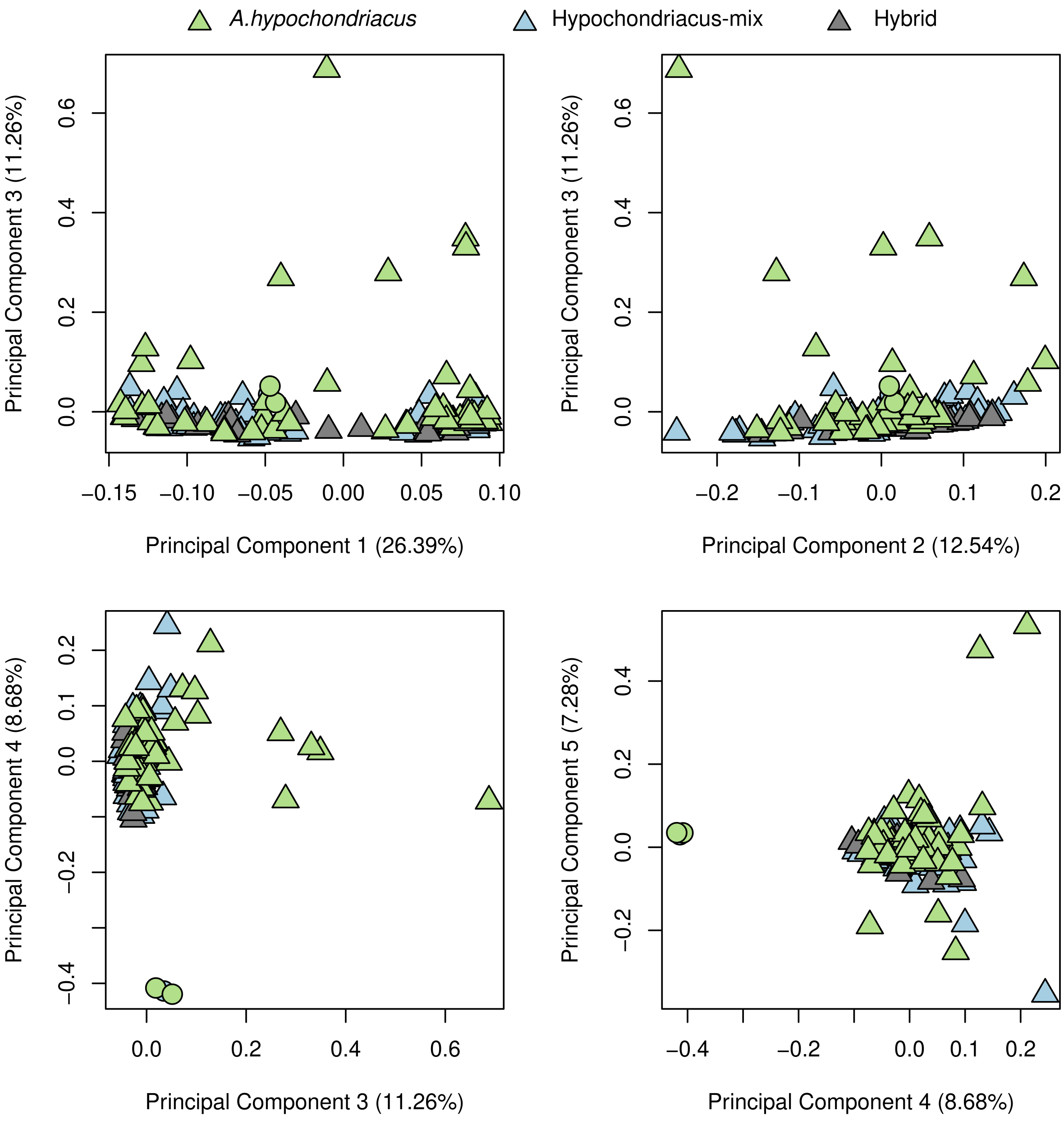


**Figure S2** PCA biplot with only *A. hypochondriacus*, Hypochondriacus-mix and Hybrid.

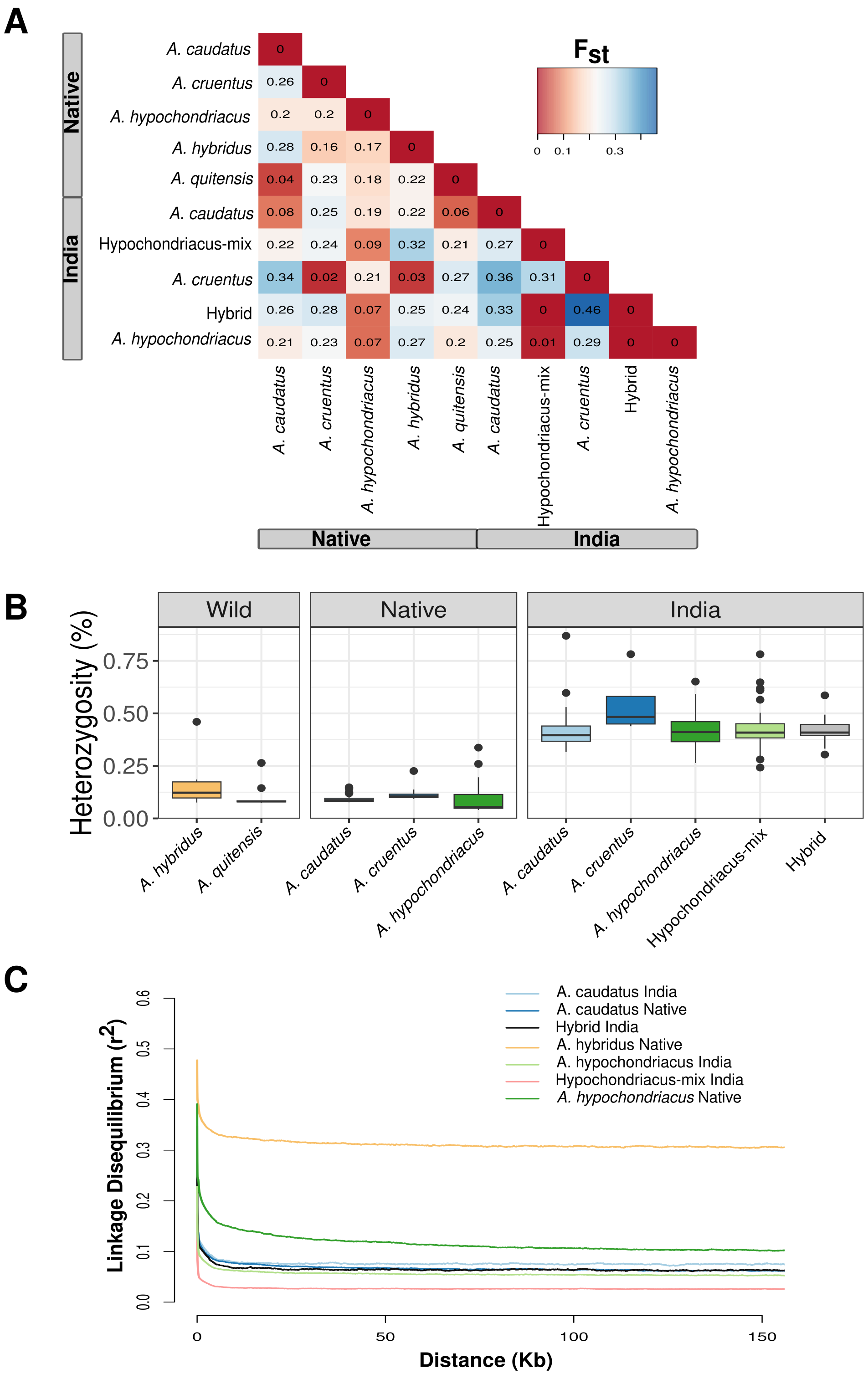


**Figure S3 Genetic diversity parameters** (A) Heatmap representing the Fst statistic between each pair of population (B) Estimate of global observed heterozygosity (in percentage) (C) Linkage disequilibrium

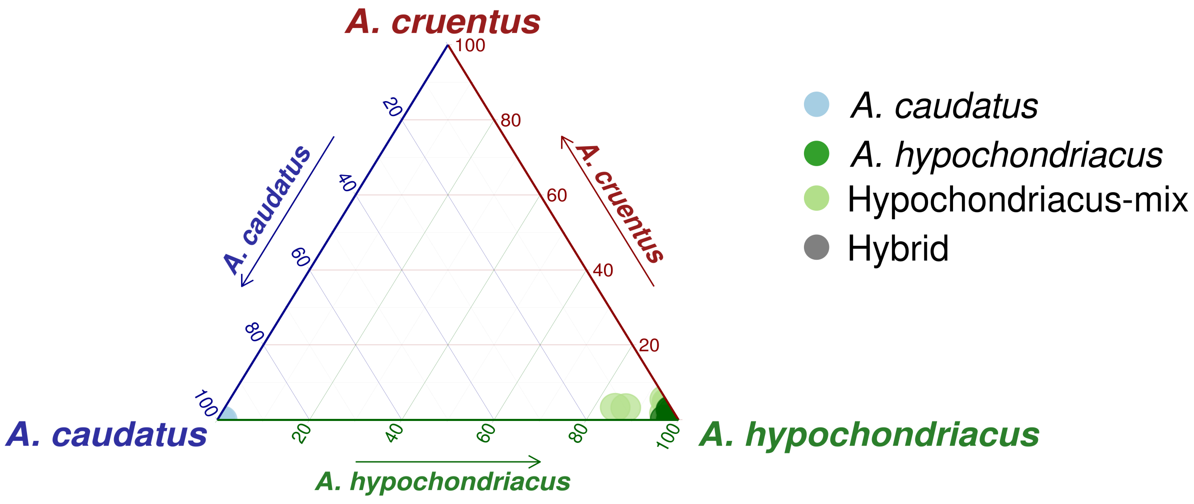


**Figure S4** Ternary plot representation for ELAI results to depict the ancestry proportion for Indian accessions using native species group as source.

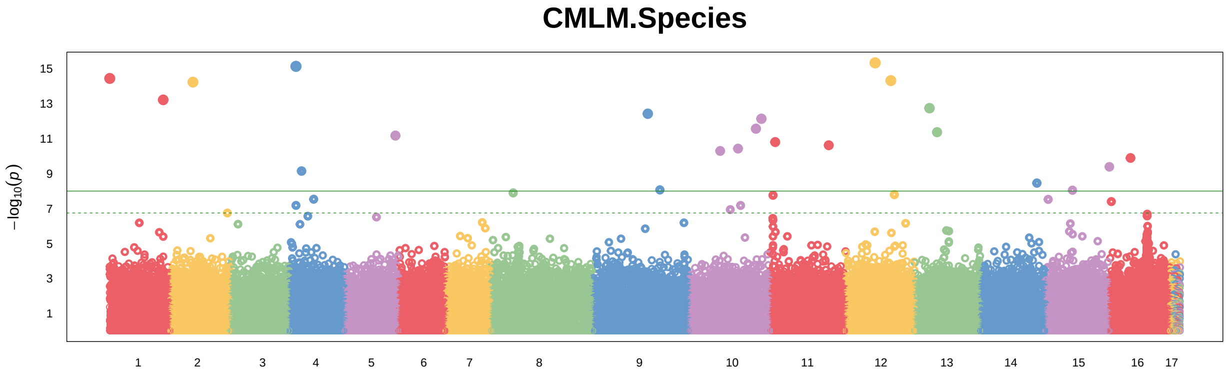


**Figure S5** Genome-wide association mapping with taxonomic characterization as phenotype using accessions of *A. caudatus* and *A. hypochondriacus* form the two range. The horizontal solid line indicates a significance level of 1E-08.

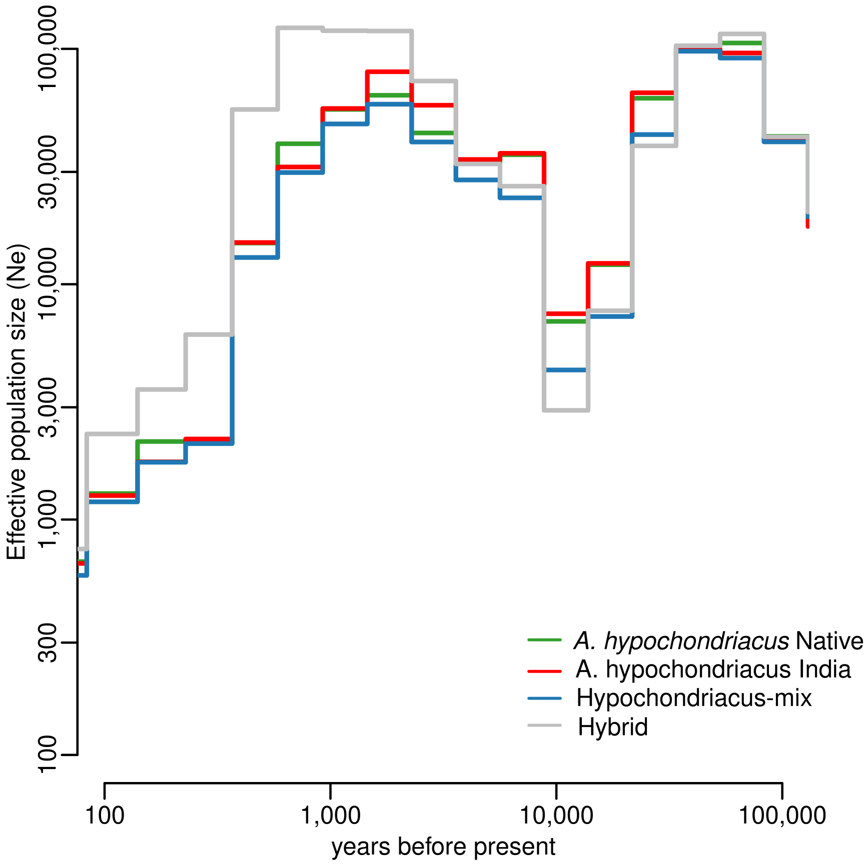


**Figure S6** Step-plot representation of effective population size estimated using popsizeABC.

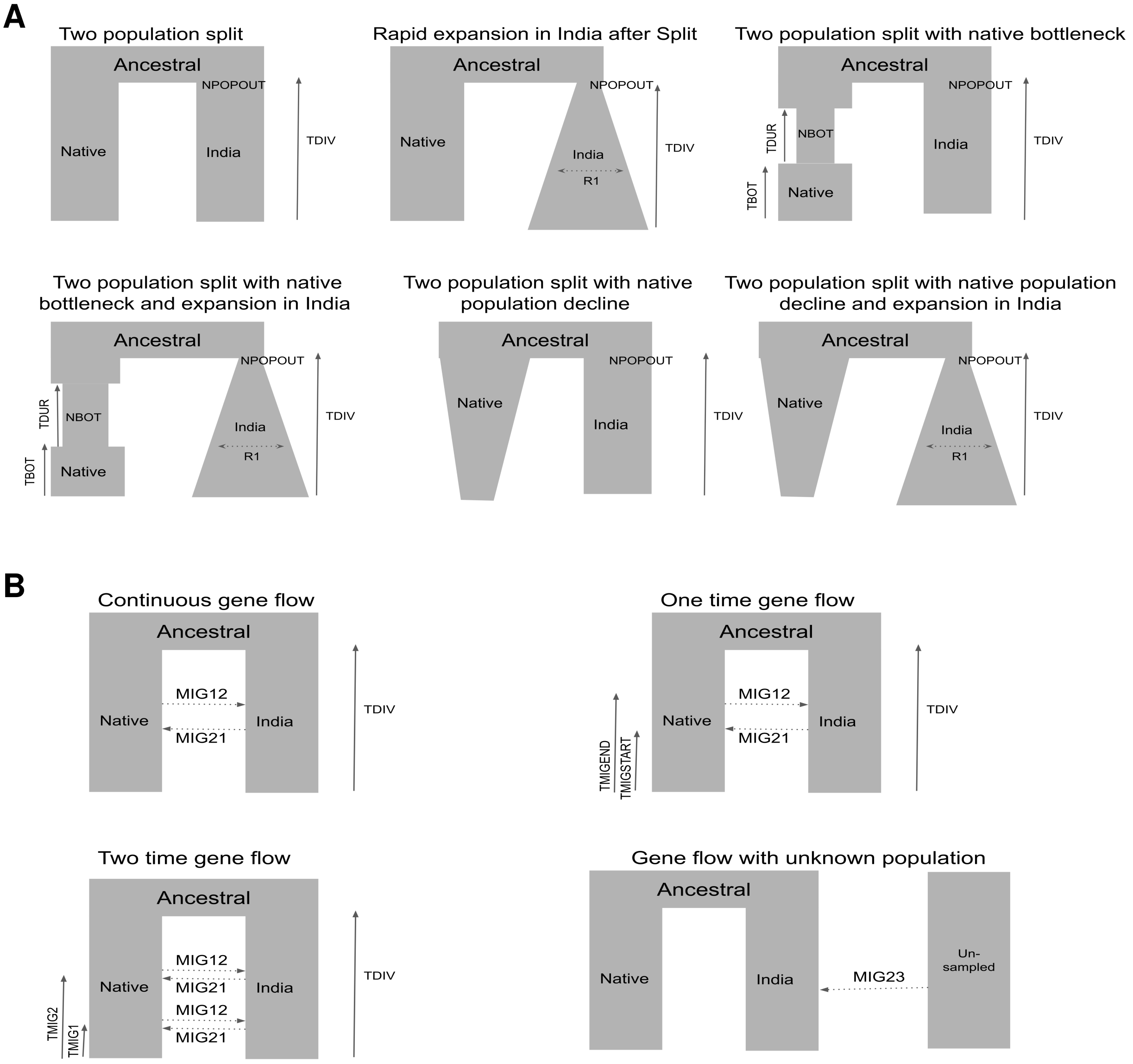


**Figure S7** Demographic models tested using fastsimcoal2 (A) Six base models, (B) Models with three gene-flow scenarios.

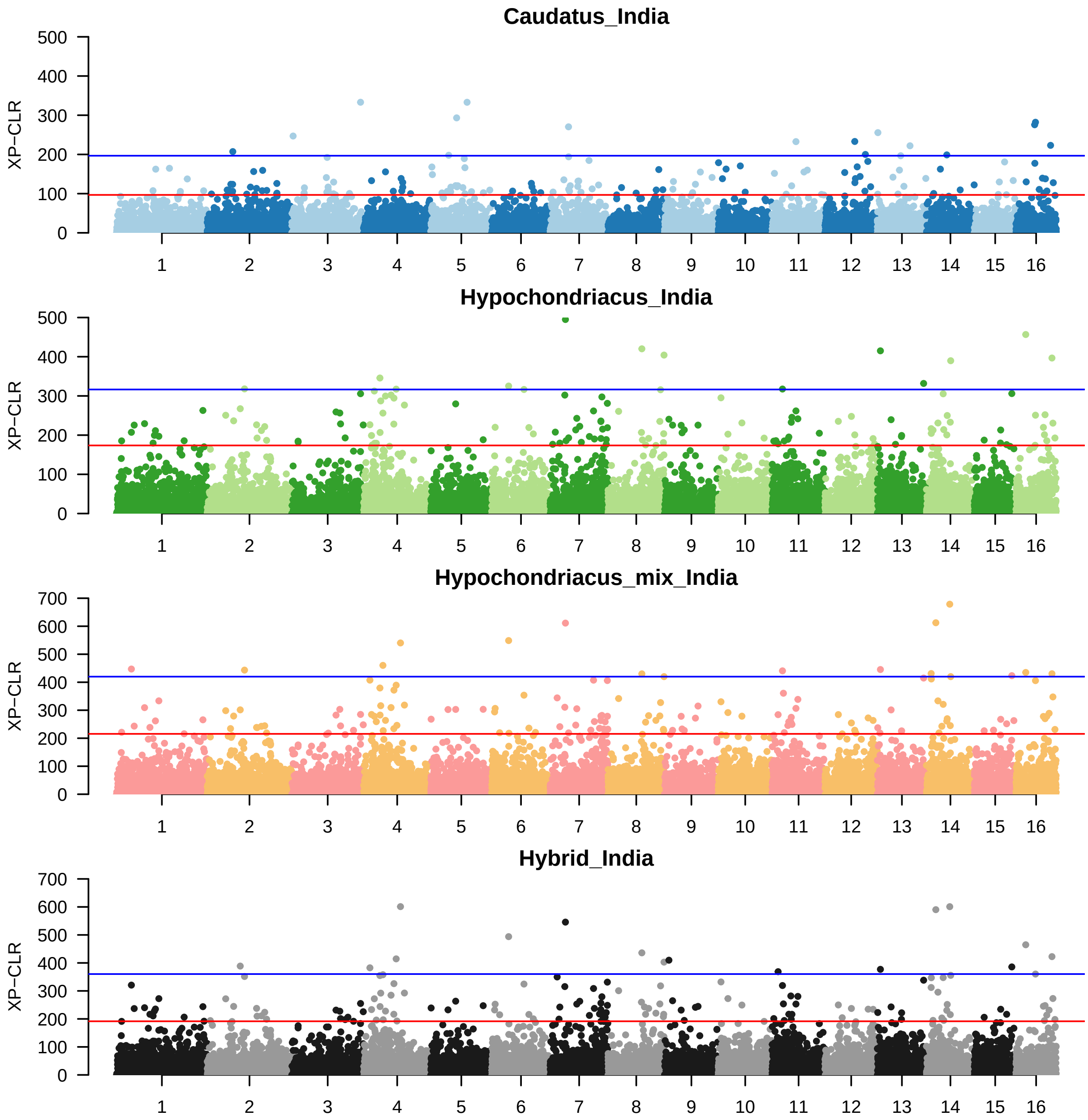


**Figure S8** Genome-wide distribution of XP-CLR statistic for two species using their native counter-part as reference population. For Hypochondriacus-mix and hybrid, *A. hypochondriacus* form native was taken as reference. Red line represent top 1 percentile and blue line represents 0.1 percentile.

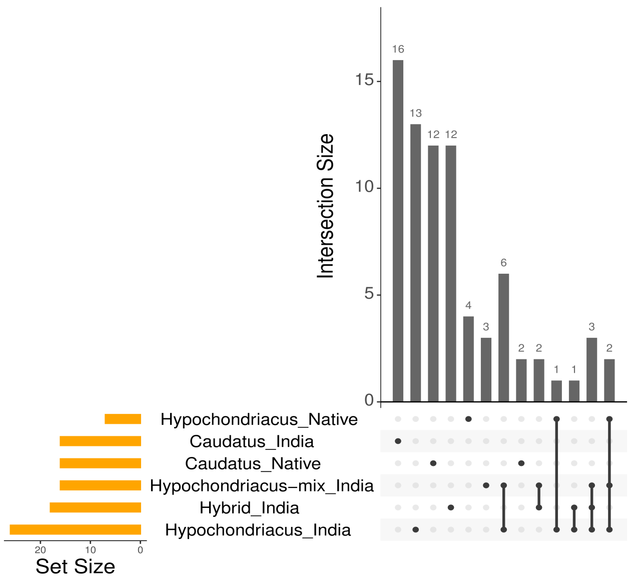


**Figure S9** Shared putative selective sweep regions identified as top 1 percentile genome-wide XP-CLR outlier with those identified per population by RAiSD

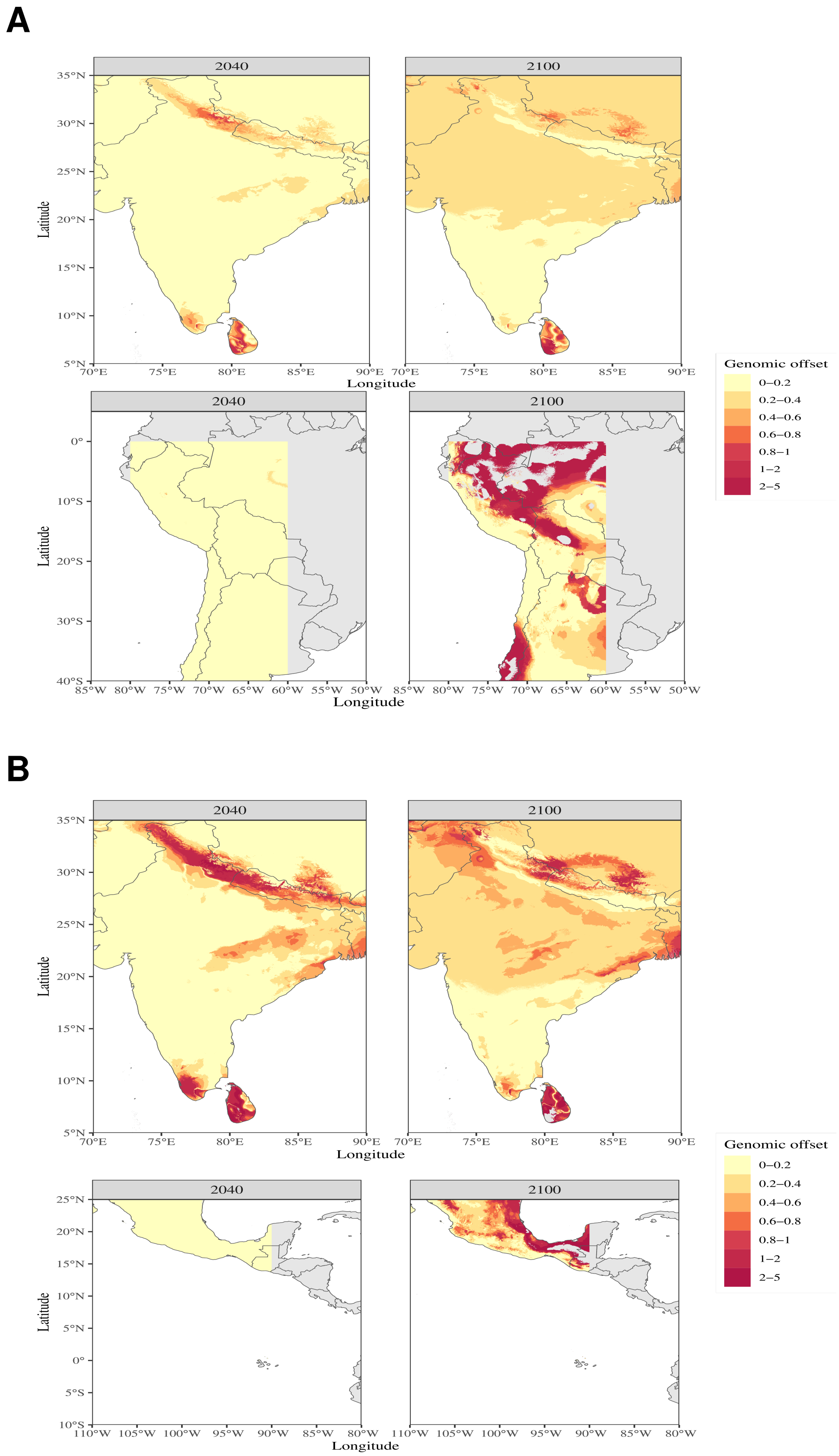


**Figure S10** Genetic offset predicted using adaptive index calculated from associated climatic variable and GEA loci in the native and introduced range for (A) *A. caudatus* and (B) *A. hypochondriacus*.

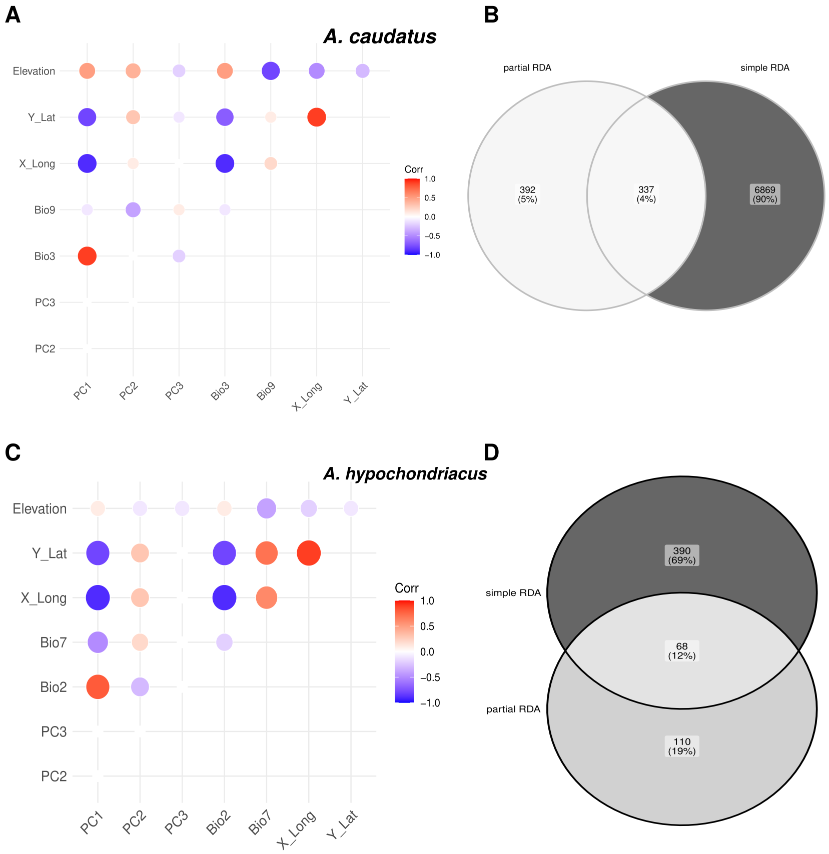


**Figure S11** Correlation among the predictor variables of RDA and overlap between constrained and unconstrained RDA analysis for the two species: (A-B) *A. caudatus* and (C-D) *A. hypochondriacus*.

### Supplementary Tables

**Table S1** List of accessions used in the study along with new assigned cluster and mapping summary.

| USDA Accession No. | Sample Name | Species (USDA) | Country Origin | Region | Mean cov. | % Aligned | Fraction SNPs missing | Included in study | New assigned cluster |
|---|---|---|---|---|---|---|---|---|---|
| PI 274278 | AM_00546 | hypochondriacus | IND | India | 8.7X | 98.5% | 9.1 | Yes | hypochondriacus_India |
| PI 480516 | AM_00547 | caudatus | IND | India | 7.6X | 98.5% | 13.3 | Yes | hypochondriacus-mix_India |
| PI 480557 | AM_00548 | hypochondriacus | IND | India | 9.5X | 98.8% | 7.3 | Yes | hypochondriacus_India |
| PI 480569 | AM_00549 | hypochondriacus | IND | India | 7.1X | 98.9% | 10.7 | Yes | hypochondriacus_India |
| PI 481091 | AM_00550 | hypochondriacus | IND | India | 10.3X | 98.8% | 6.9 | Yes | hypochondriacus_India |
| PI 481184 | AM_00551 | hypochondriacus | IND | India | 8.4X | 98.6% | 11.1 | Yes | hypochondriacus_India |
| PI 481336 | AM_00552 | hypochondriacus | IND | India | 6.7X | 98.6% | 17.2 | Yes | hypochondriacus_India |
| PI 669884 | AM_00553 | hypochondriacus | IND | India | 8.8X | 98.4% | 11 | Yes | hypochondriacus_India |
| PI 669931 | AM_00554 | hypochondriacus | IND | India | 4.6X | 98.6% | 27.2 | Yes | hypochondriacus_India |
| PI 566897 | AM_00558 | cruentus | IND | India | 7.6X | 98.1% | 18.3 | Yes | cruentus_India |
| Ames 15204 | AM_00601 | hypochondriacus | IND | India | 4.1X | 99.1% | 25.2 | Yes | hypochondriacus_India |
| Ames 15210 | AM_00602 | hypochondriacus | IND | India | 1.1X | 99.3% | 64.3 | Yes | hypochondriacus_India |
| Ames 15301 | AM_00604 | hypochondriacus | IND | India | 9.8X | 98.4% | 8.2 | Yes | hypochondriacus_India |
| Ames 2257 | AM_00607 | hypochondriacus | IND | India | 6.2X | 98.6% | 16.7 | Yes | hypochondriacus_India |
| Ames 5138 | AM_00609 | hypochondriacus | IND | India | 10.0X | 98.7% | 7.4 | Yes | hypochondriacus_India |
| Ames 5145 | AM_00610 | hypochondriacus | IND | India | 8.8X | 98.7% | 9.3 | Yes | hypochondriacus_India |
| Ames 5386 | AM_00613 | cruentus | IND | India | 8.5X | 98.1% | 19 | Yes | cruentus_India |
| Ames 5585 | AM_00637 | hypochondriacus | IND | India | 10.0X | 98.4% | 7.8 | Yes | hypochondriacus_India |
| Ames 5586 | AM_00638 | hypochondriacus | IND | India | 10.2X | 98.4% | 8.5 | Yes | hypochondriacus_India |
| Ames 5587 | AM_00639 | hypochondriacus | IND | India | 7.4X | 98.4% | 13.8 | Yes | hypochondriacus_India |
| Ames 5588 | AM_00640 | hypochondriacus | IND | India | 9.1X | 98.4% | 7.9 | Yes | hypochondriacus_India |
| Ames 5593 | AM_00643 | hypochondriacus | IND | India | 10.7X | 98.6% | 6.1 | Yes | hypochondriacus_India |
| Ames 5603 | AM_00644 | hypochondriacus | IND | India | 5.1X | 99.2% | 18.3 | Yes | hypochondriacus_India |
| Ames 5608 | AM_00645 | hypochondriacus | IND | India | 9.8X | 98.6% | 7.1 | Yes | hypochondriacus_India |
| Ames 5611 | AM_00646 | hypochondriacus | IND | India | 8.1X | 98.3% | 12.4 | Yes | hypochondriacus_India |
| Ames 5612 | AM_00647 | hypochondriacus | IND | India | 10.0X | 98.6% | 7.6 | Yes | hypochondriacus_India |
| Ames 5613 | AM_00648 | hypochondriacus | IND | India | 3.9X | 98.9% | 31.2 | Yes | hypochondriacus_India |
| PI 166045 | AM_00664 | caudatus | IND | India | 7.9X | 98.3% | 19.6 | Yes | caudatus_India |
| PI 166107 | AM_00665 | caudatus | IND | India | 8.3X | 98.0% | 20.8 | Yes | caudatus_India |
| PI 175039 | AM_00666 | caudatus | IND | India | 9.4X | 98.2% | 17.2 | Yes | caudatus_India |
| PI 175040 | AM_00667 | hypochondriacus | IND | India | 7.5X | 98.7% | 16.7 | Yes | hypochondriacus_India |
| PI 274277 | AM_00668 | hypochondriacus | IND | India | 9.6X | 98.3% | 8.7 | Yes | hypochondriacus_India |
| PI 274278 | AM_00669 | hypochondriacus | IND | India | 6.6X | 98.5% | 16.3 | Yes | hypochondriacus_India |
| PI 274279 | AM_00670 | hypochondriacus | IND | India | 8.0X | 97.9% | 12.1 | Yes | hypochondriacus_India |
| PI 288278 | AM_00671 | cruentus | IND | India | 7.8X | 98.4% | 18.9 | Yes | cruentus_India |

| | | | | | | | | | |
|---|---|---|---|---|---|---|---|---|---|
| PI 288279 | AM_00672 | caudatus | IND | India | 8.7X | 98.2% | 18 | Yes | caudatus_India |
| PI 480487 | AM_00675 | caudatus | IND | India | 8.4X | 98.6% | 9.6 | Yes | hypochondriacus-mix_India |
| PI 480491 | AM_00676 | caudatus | IND | India | 11.1X | 99.0% | 4.7 | Yes | hypochondriacus-mix_India |
| PI 480493 | AM_00677 | caudatus | IND | India | 0.9X | 98.9% | 59.6 | Yes | hypochondriacus-mix_India |
| PI 480494 | AM_00678 | caudatus | IND | India | 7.4X | 98.6% | 12.3 | Yes | hypochondriacus-mix_India |
| PI 480498 | AM_00679 | caudatus | IND | India | 8.3X | 98.6% | 10.3 | Yes | hypochondriacus-mix_India |
| PI 480500 | AM_00680 | caudatus | IND | India | 6.7X | 98.7% | 14.4 | Yes | hypochondriacus-mix_India |
| PI 480503 | AM_00681 | caudatus | IND | India | 8.3X | 98.7% | 15.4 | Yes | hypochondriacus-mix_India |
| PI 480514 | AM_00682 | caudatus | IND | India | 10.5X | 98.4% | 13.9 | Yes | caudatus_India |
| PI 480516 | AM_00683 | caudatus | IND | India | 8.1X | 99.1% | 8.4 | Yes | hypochondriacus-mix_India |
| PI 480525 | AM_00684 | caudatus | IND | India | 6.0X | 99.0% | 17.9 | Yes | hypochondriacus-mix_India |
| PI 480546 | AM_00685 | caudatus | IND | India | 8.6X | 98.4% | 19.4 | Yes | caudatus_India |
| PI 480568 | AM_00686 | caudatus | IND | India | 7.2X | 98.6% | 14 | Yes | hypochondriacus-mix_India |
| PI 480574 | AM_00687 | caudatus | IND | India | 9.6X | 99.1% | 6.3 | Yes | hypochondriacus-mix_India |
| PI 480575 | AM_00688 | caudatus | IND | India | 4.2X | 98.9% | 30.6 | Yes | hypochondriacus-mix_India |
| PI 480576 | AM_00689 | caudatus | IND | India | 1.0X | 98.6% | 63.6 | Yes | caudatus_India |
| PI 480577 | AM_00690 | caudatus | IND | India | 7.0X | 98.6% | 15.7 | Yes | hypochondriacus-mix_India |
| PI 480591 | AM_00691 | caudatus | IND | India | 5.9X | 98.8% | 17.1 | Yes | hypochondriacus-mix_India |
| PI 480596 | AM_00692 | caudatus | IND | India | 10.0X | 98.6% | 5.3 | Yes | hypochondriacus-mix_India |
| PI 480616 | AM_00693 | caudatus | IND | India | 7.3X | 98.2% | 19 | Yes | caudatus_India |
| PI 480623 | AM_00694 | caudatus | IND | India | 7.8X | 98.8% | 12.1 | Yes | hypochondriacus-mix_India |
| PI 480655 | AM_00695 | caudatus | IND | India | 8.9X | 99.0% | 6.7 | Yes | hypochondriacus-mix_India |
| PI 480656 | AM_00696 | caudatus | IND | India | 1.6X | 96.9% | 42.7 | Yes | hypochondriacus-mix_India |
| PI 480659 | AM_00697 | caudatus | IND | India | 7.2X | 98.9% | 10.1 | Yes | hypochondriacus-mix_India |
| PI 480660 | AM_00698 | caudatus | IND | India | 7.6X | 98.9% | 10.8 | Yes | hypochondriacus-mix_India |
| PI 480666 | AM_00699 | caudatus | IND | India | 1.6X | 96.8% | 41.4 | Yes | hypochondriacus-mix_India |
| PI 480674 | AM_00700 | caudatus | IND | India | 5.6X | 98.6% | 17.7 | Yes | hypochondriacus-mix_India |

| | | | | | | | | | |
|---|---|---|---|---|---|---|---|---|---|
| PI 480675 | AM_00701 | caudatus | IND | India | 1.8X | 98.6% | 38.7 | Yes | hypochondriacus-mix_India |
| PI 480676 | AM_00702 | caudatus | IND | India | 10.1X | 98.5% | 5.6 | Yes | hypochondriacus-mix_India |
| PI 480677 | AM_00703 | caudatus | IND | India | 7.8X | 98.5% | 9.2 | Yes | hypochondriacus-mix_India |
| PI 480679 | AM_00704 | hybr. | IND | India | 6.9X | 99.0% | 9.8 | Yes | hybrid_India |
| PI 480742 | AM_00705 | caudatus | IND | India | 7.5X | 98.6% | 11.9 | Yes | hypochondriacus-mix_India |
| PI 480748 | AM_00706 | caudatus | IND | India | 1.6X | 99.2% | 43.5 | Yes | hypochondriacus-mix_India |
| PI 480749 | AM_00707 | caudatus | IND | India | 0.8X | 93.3% | 64.6 | Yes | hypochondriacus-mix_India |
| PI 480750 | AM_00708 | caudatus | IND | India | 2.4X | 99.3% | 39.4 | Yes | hypochondriacus-mix_India |
| PI 480760 | AM_00709 | caudatus | IND | India | 8.3X | 98.9% | 7.5 | Yes | hypochondriacus-mix_India |
| PI 480762 | AM_00710 | hybr. | IND | India | 5.9X | 98.7% | 19.4 | Yes | hybrid_India |
| PI 480767 | AM_00711 | hybr. | IND | India | 6.8X | 98.6% | 14.9 | Yes | hybrid_India |
| PI 480788 | AM_00712 | caudatus | IND | India | 1.5X | 99.0% | 44.4 | Yes | hypochondriacus-mix_India |
| PI 480797 | AM_00713 | hybr. | IND | India | 8.6X | 98.7% | 8 | Yes | hybrid_India |
| PI 480804 | AM_00714 | caudatus | IND | India | 0.9X | 97.1% | 64 | Yes | hypochondriacus-mix_India |
| PI 480805 | AM_00715 | caudatus | IND | India | 8.6X | 98.9% | 7 | Yes | hypochondriacus-mix_India |
| PI 480816 | AM_00716 | caudatus | IND | India | 1.6X | 99.1% | 43.1 | Yes | hypochondriacus-mix_India |
| PI 480820 | AM_00717 | hybr. | IND | India | 3.8X | 99.4% | 25.2 | Yes | hybrid_India |
| PI 480822 | AM_00718 | caudatus | IND | India | 0.9X | 99.4% | 63.1 | Yes | hypochondriacus-mix_India |
| PI 480826 | AM_00719 | hybr. | IND | India | 8.6X | 99.0% | 6.5 | Yes | hybrid_India |
| PI 480827 | AM_00720 | hybr. | IND | India | 5.0X | 98.6% | 16.5 | Yes | hybrid_India |
| PI 480828 | AM_00721 | caudatus | IND | India | 5.1X | 98.3% | 20.4 | Yes | hypochondriacus-mix_India |
| PI 480833 | AM_00722 | caudatus | IND | India | 6.3X | 98.4% | 15.6 | Yes | hypochondriacus-mix_India |
| PI 480834 | AM_00723 | hybr. | IND | India | 9.4X | 98.5% | 6.5 | Yes | hybrid_India |
| PI 480836 | AM_00724 | caudatus | IND | India | 8.8X | 99.0% | 7.1 | Yes | hypochondriacus-mix_India |
| PI 480837 | AM_00725 | hybr. | IND | India | 6.4X | 98.5% | 14.7 | Yes | hybrid_India |
| PI 480845 | AM_00726 | hybr. | IND | India | 4.9X | 98.7% | 17 | Yes | hybrid_India |
| PI 480848 | AM_00727 | caudatus | IND | India | 2.1X | 99.1% | 33.9 | Yes | hypochondriacus-mix_India |
| PI 480851 | AM_00728 | hybr. | IND | India | 10.5X | 99.0% | 5 | Yes | hybrid_India |
| PI 480854 | AM_00729 | caudatus | IND | India | 8.4X | 98.6% | 7.6 | Yes | hypochondriacus-mix_India |

| | | | | | | | | | |
|---|---|---|---|---|---|---|---|---|---|
| PI 480855 | AM_00730 | caudatus | IND | India | 2.2X | 98.6% | 31.7 | Yes | hypochondriacus-mix_India |
| PI 480861 | AM_00732 | hybr. | IND | India | 1.0X | 98.8% | 56.8 | Yes | hybrid_India |
| PI 480864 | AM_00733 | caudatus | IND | India | 0.3X | 98.9% | 84.6 | No | NA |
| PI 480902 | AM_00734 | hybr. | IND | India | 1.0X | 98.9% | 58.1 | Yes | hybrid_India |
| PI 480923 | AM_00735 | hybr. | IND | India | 1.3X | 98.7% | 47.9 | Yes | hybrid_India |
| PI 480993 | AM_00736 | caudatus | IND | India | 0.5X | 99.0% | 75.9 | Yes | hypochondriacus-mix_India |
| PI 480994 | AM_00737 | caudatus | IND | India | 7.6X | 98.8% | 8.3 | Yes | hypochondriacus-mix_India |
| PI 480999 | AM_00738 | caudatus | IND | India | 7.5X | 99.1% | 9.4 | Yes | hypochondriacus-mix_India |
| PI 481000 | AM_00739 | caudatus | IND | India | 0.4X | 98.9% | 78.4 | Yes | hypochondriacus-mix_India |
| PI 481002 | AM_00740 | hybr. | IND | India | 0.8X | 98.6% | 62.6 | Yes | hybrid_India |
| PI 481004 | AM_00741 | caudatus | IND | India | 0.9X | 88.9% | 58.1 | Yes | hypochondriacus-mix_India |
| PI 481005 | AM_00742 | hybr. | IND | India | 0.5X | 98.9% | 76.9 | Yes | hybrid_India |
| PI 481006 | AM_00743 | caudatus | IND | India | 0.5X | 99.1% | 74.5 | Yes | hypochondriacus-mix_India |
| PI 481007 | AM_00744 | caudatus | IND | India | 1.0X | 99.0% | 55.5 | Yes | hypochondriacus-mix_India |
| PI 481012 | AM_00745 | hybr. | IND | India | 0.2X | 98.5% | 88.7 | No | NA |
| PI 481019 | AM_00746 | hybr. | IND | India | 9.2X | 98.8% | 8.4 | Yes | hybrid_India |
| PI 481020 | AM_00747 | caudatus | IND | India | 1.5X | 98.7% | 47.8 | Yes | hypochondriacus-mix_India |
| PI 481022 | AM_00748 | hybr. | IND | India | 1.7X | 98.9% | 41.3 | Yes | hybrid_India |
| PI 481024 | AM_00749 | hybr. | IND | India | 1.3X | 98.8% | 48.5 | Yes | hybrid_India |
| PI 481025 | AM_00750 | caudatus | IND | India | 0.6X | 99.0% | 71.6 | Yes | hypochondriacus-mix_India |
| PI 481026 | AM_00751 | caudatus | IND | India | 0.8X | 99.0% | 62.6 | Yes | hypochondriacus-mix_India |
| PI 481028 | AM_00752 | caudatus | IND | India | 0.3X | 98.8% | 83.4 | No | NA |
| PI 481030 | AM_00753 | hybr. | IND | India | 1.7X | 98.0% | 42.4 | Yes | hybrid_India |
| PI 481032 | AM_00754 | caudatus | IND | India | 8.5X | 98.5% | 7.6 | Yes | hypochondriacus-mix_India |
| PI 481033 | AM_00755 | hybr. | IND | India | 0.6X | 98.9% | 71.3 | Yes | hybrid_India |
| PI 481037 | AM_00756 | caudatus | IND | India | 0.5X | 98.4% | 77.3 | Yes | caudatus_India |
| PI 481040 | AM_00757 | caudatus | IND | India | 1.4X | 98.3% | 44.4 | Yes | hypochondriacus-mix_India |
| PI 481041 | AM_00758 | caudatus | IND | India | 8.2X | 98.8% | 13.1 | Yes | hypochondriacus-mix_India |
| PI 481042 | AM_00759 | hypochondriacus | IND | India | 1.9X | 99.2% | 36 | Yes | hypochondriacus_India |
| PI 481043 | AM_00760 | caudatus | IND | India | 0.6X | 98.6% | 72.3 | Yes | hypochondriacus-mix_India |
| PI 481045 | AM_00762 | caudatus | IND | India | 1.0X | 99.1% | 58.6 | Yes | hypochondriacus-mix_India |

| | | | | | | | | | |
|---|---|---|---|---|---|---|---|---|---|
| PI 481046 | AM_00763 | caudatus | IND | India | 0.7X | 98.2% | 70.5 | Yes | caudatus_India |
| PI 481047 | AM_00764 | caudatus | IND | India | 4.0X | 98.7% | 25.1 | Yes | caudatus_India |
| PI 481048 | AM_00765 | caudatus | IND | India | 8.3X | 98.2% | 12.4 | Yes | hypochondriacus-mix_India |
| PI 481051 | AM_00766 | caudatus | IND | India | 0.3X | 98.1% | 86.8 | No | NA |
| PI 481052 | AM_00767 | caudatus | IND | India | 2.7X | 98.7% | 25.5 | Yes | hypochondriacus-mix_India |
| PI 481053 | AM_00768 | caudatus | IND | India | 0.9X | 99.1% | 58.9 | Yes | hypochondriacus-mix_India |
| PI 481054 | AM_00769 | caudatus | IND | India | 0.7X | 98.7% | 68.3 | Yes | hypochondriacus-mix_India |
| PI 481056 | AM_00770 | hybr. | IND | India | 0.8X | 98.8% | 64.8 | Yes | hybrid_India |
| PI 481057 | AM_00771 | caudatus | IND | India | 0.4X | 98.2% | 82 | No | NA |
| PI 481058 | AM_00772 | caudatus | IND | India | 5.3X | 98.7% | 23.5 | Yes | hypochondriacus-mix_India |
| PI 481059 | AM_00773 | caudatus | IND | India | 2.1X | 98.7% | 49.2 | Yes | hypochondriacus-mix_India |
| PI 481061 | AM_00774 | caudatus | IND | India | 0.9X | 98.8% | 62.6 | Yes | hypochondriacus-mix_India |
| PI 481064 | AM_00775 | caudatus | IND | India | 10.1X | 98.6% | 15.4 | Yes | caudatus_India |
| PI 481066 | AM_00776 | caudatus | IND | India | 1.3X | 98.6% | 51.7 | Yes | hypochondriacus-mix_India |
| PI 481067 | AM_00777 | caudatus | IND | India | 1.0X | 98.9% | 62.4 | Yes | hypochondriacus-mix_India |
| PI 481072 | AM_00778 | caudatus | IND | India | 8.9X | 98.0% | 6.8 | Yes | hypochondriacus-mix_India |
| PI 481073 | AM_00779 | hybr. | IND | India | 1.2X | 98.5% | 55.9 | Yes | hybrid_India |
| PI 481074 | AM_00780 | hypochondriacus | IND | India | 1.2X | 98.5% | 54 | Yes | hypochondriacus_India |
| PI 481075 | AM_00781 | hypochondriacus | IND | India | 0.9X | 98.8% | 59.6 | Yes | hypochondriacus_India |
| PI 481076 | AM_00782 | hypochondriacus | IND | India | 1.5X | 98.9% | 44.2 | Yes | hypochondriacus_India |
| PI 481087 | AM_00783 | caudatus | IND | India | 0.3X | 98.4% | 84.2 | No | NA |
| PI 481090 | AM_00784 | hybr. | IND | India | 7.8X | 99.2% | 11.3 | Yes | hybrid_India |
| PI 481091 | AM_00785 | hypochondriacus | IND | India | 0.6X | 99.2% | 71.6 | Yes | hypochondriacus_India |
| PI 481092 | AM_00786 | hybr. | IND | India | 6.1X | 98.7% | 19.6 | Yes | hybrid_India |
| PI 481093 | AM_00787 | hypochondriacus | IND | India | 0.4X | 99.3% | 77.4 | Yes | hypochondriacus_India |
| PI 481098 | AM_00788 | caudatus | IND | India | 0.3X | 98.7% | 87.1 | No | NA |
| PI 481101 | AM_00789 | caudatus | IND | India | 0.5X | 98.6% | 75.4 | Yes | caudatus_India |
| PI 481106 | AM_00790 | hypochondriacus | IND | India | 0.6X | 99.2% | 71.8 | Yes | hypochondriacus_India |
| PI 481107 | AM_00791 | hypochondriacus | IND | India | 0.4X | 99.1% | 77.5 | Yes | hypochondriacus_India |
| PI 481108 | AM_00792 | hypochondriacus | IND | India | 0.5X | 98.8% | 75.2 | Yes | hypochondriacus_India |
| PI 481109 | AM_00793 | hypochondriacus | IND | India | 0.5X | 98.7% | 76.1 | Yes | hypochondriacus_India |
| PI 481110 | AM_00794 | hypochondriacus | IND | India | 1.9X | 99.0% | 38 | Yes | hypochondriacus_India |
| PI 481113 | AM_00795 | caudatus | IND | India | 1.6X | 98.8% | 45.8 | Yes | hypochondriacus-mix_India |
| PI 481116 | AM_00796 | caudatus | IND | India | 1.3X | 98.6% | 49.1 | Yes | hypochondriacus-mix_India |

| | | | | | | | | | |
|---|---|---|---|---|---|---|---|---|---|
| PI 481117 | AM_00797 | caudatus | IND | India | 9.0X | 98.3% | 14.3 | Yes | caudatus_India |
| PI 481124 | AM_00798 | caudatus | IND | India | 0.4X | 99.0% | 78.3 | Yes | hypochondriacus-mix_India |
| PI 481125 | AM_00799 | caudatus | IND | India | 0.7X | 98.9% | 64.7 | Yes | hypochondriacus-mix_India |
| PI 481374 | AM_00800 | caudatus | IND | India | 1.7X | 95.0% | 45.9 | Yes | caudatus_India |
| PI 615696 | AM_00864 | hypochondriacus | IND | India | 1.6X | 98.2% | 46.3 | Yes | hypochondriacus_India |
| PI 619236 | AM_00865 | caudatus | IND | India | 2.8X | 98.0% | 31.8 | Yes | caudatus_India |
| PI 636183 | AM_00870 | hypochondriacus | IND | India | 0.8X | 98.7% | 65.2 | Yes | hypochondriacus_India |
| PI 636184 | AM_00871 | hypochondriacus | IND | India | 0.3X | 98.8% | 86.2 | No | NA |
| PI 636185 | AM_00872 | hypochondriacus | IND | India | 8.0X | 98.7% | 8.9 | Yes | hypochondriacus_India |
| PI 636186 | AM_00873 | hypochondriacus | IND | India | 0.6X | 99.1% | 72 | Yes | hypochondriacus_India |
| PI 636187 | AM_00874 | hypochondriacus | IND | India | 0.4X | 98.8% | 80.9 | No | NA |
| PI 636188 | AM_00875 | hypochondriacus | IND | India | 0.3X | 98.3% | 81.5 | No | NA |
| PI 636190 | AM_00876 | hypochondriacus | IND | India | 1.8X | 96.0% | 39.7 | Yes | hypochondriacus_India |
| PI 636191 | AM_00877 | hypochondriacus | IND | India | 1.9X | 98.9% | 40.7 | Yes | hypochondriacus_India |
| PI 636192 | AM_00878 | hypochondriacus | IND | India | 1.3X | 98.3% | 50.9 | Yes | hypochondriacus_India |
| PI 636193 | AM_00879 | caudatus | IND | India | 0.7X | 98.4% | 71.1 | Yes | caudatus_India |
| PI 669855 | AM_00987 | caudatus | IND | India | 5.6X | 93.8% | 26.6 | Yes | caudatus_India |
| PI 669883 | AM_00988 | caudatus | IND | India | 0.6X | 98.7% | 73.5 | Yes | caudatus_India |
| PI 669885 | AM_00989 | caudatus | IND | India | 0.5X | 98.7% | 77.5 | Yes | caudatus_India |
| PI 669888 | AM_00990 | caudatus | IND | India | 0.7X | 98.5% | 71.5 | Yes | caudatus_India |
| PI 669889 | AM_00991 | caudatus | IND | India | 0.1X | 98.3% | 92.9 | No | NA |
| PI 669891 | AM_00992 | caudatus | IND | India | 0.8X | 98.7% | 70.2 | Yes | caudatus_India |
| PI 669895 | AM_00993 | caudatus | IND | India | 0.9X | 99.0% | 62.9 | Yes | hypochondriacus-mix_India |
| PI 669901 | AM_00994 | caudatus | IND | India | 3.3X | 98.4% | 37 | Yes | caudatus_India |
| PI 669908 | AM_00995 | caudatus | IND | India | 4.4X | 98.5% | 33.5 | Yes | hypochondriacus-mix_India |
| PI 669930 | AM_00996 | caudatus | IND | India | 1.2X | 97.9% | 60.3 | Yes | caudatus_India |
| PI 669934 | AM_00997 | caudatus | IND | India | 0.0X | 97.7% | NA | No | NA |
| PI 674256 | AM_01009 | hypochondriacus | IND | India | 1.7X | 97.7% | 42.1 | Yes | hypochondriacus_India |
| PI 689732 | AM_01021 | caudatus | IND | India | 1.5X | 98.4% | 53.7 | Yes | caudatus_India |
| PI 689735 | AM_01023 | hypochondriacus | IND | India | 1.1X | 98.1% | 61.3 | Yes | hypochondriacus_India |
| PI 689736 | AM_01024 | hypochondriacus | IND | India | 0.2X | 98.0% | 88.5 | No | NA |
| PI 689738 | AM_01025 | hypochondriacus | IND | India | 0.4X | 95.7% | 79.5 | Yes | hypochondriacus_India |
| PI 566897 | AM_01059 | cruentus | IND | India | 0.8X | 97.3% | 67.4 | Yes | cruentus_India |
| AMA132 | AM_01094 | cruentus | IND | India | 0.7X | 99.0% | 70.7 | Yes | hypochondriacus_India |
| AMA125 | AMA125 | caudatus | PER | Native | 12.4X | 97.5% | 9.9 | Yes | caudatus_Native |
| AMA155 | AMA155 | cruentus | USA | Native | 9.6X | 91.1% | 9.9 | Yes | cruentus_Native |
| Ames 2085 | Ames2085 | hypochondriacus | MEX | Native | 11.1X | 97.5% | 3.2 | Yes | hypochondriacus_Native |
| Ames 5232 | Ames5232 | hybridus | PER | Native | 8.3X | 95.5% | 19.9 | Yes | hybridus_SA_Native |
| Ames 5247 | Ames5247 | quitensis | ECU | Native | 10.4X | 97.3% | 10.2 | Yes | quitensis_Native |

| | | | | | | | | | |
|---|---|---|---|---|---|---|---|---|---|
| Ames 5302 | Ames5302 | caudatus | PER | Native | 10.0X | 95.8% | 10.8 | Yes | caudatus_Native |
| Ames 5342 | Ames5342 | quitensis | PER | Native | 9.4X | 97.7% | 9.6 | Yes | quitensis_Native |
| Ames 5457 | Ames5457 | hypochondriacus | MEX | Native | 12.1X | 98.0% | 3.6 | Yes | hypochondriacus_Native |
| Ames 5552 | Ames5552 | cruentus | MEX | Native | 9.3X | 97.1% | 10.3 | Yes | cruentus_Native |
| PI 433228 | PI433228 | cruentus | GTM | Native | 8.0X | 97.5% | 10.7 | Yes | cruentus_Native |
| PI 451826 | PI451826 | cruentus | GTM | Native | 15.8X | 93.6% | 8.8 | Yes | cruentus_Native |
| PI 481957 | PI481957 | caudatus | PER | Native | 11.2X | 97.2% | 9.8 | Yes | caudatus_Native |
| PI 481960 | PI481960 | caudatus | PER | Native | 12.7X | 97.4% | 9.4 | Yes | caudatus_Native |
| PI 481965 | PI481965 | caudatus | PER | Native | 11.9X | 95.1% | 9.9 | Yes | caudatus_Native |
| PI 490431 | PI490431 | caudatus | PER | Native | 11.0X | 96.7% | 10.3 | Yes | caudatus_Native |
| PI 490459 | PI490459 | caudatus | BOL | Native | 12.7X | 96.2% | 9.8 | Yes | caudatus_Native |
| PI 490466 | PI490466 | quitensis | PER | Native | 11.8X | 97.4% | 10 | Yes | quitensis_Native |
| PI 490489 | PI490489 | hybridus | PER | Native | 12.1X | 95.3% | 18.2 | Yes | hybridus_CA_Native |
| PI 490518 | PI490518 | caudatus | PER | Native | 11.7X | 96.7% | 9.7 | Yes | caudatus_Native |
| PI 490561 | PI490561 | caudatus | PER | Native | 9.3X | 94.4% | 11.2 | Yes | caudatus_Native |
| PI 490604 | PI490604 | caudatus | BOL | Native | 8.0X | 96.0% | 11.6 | Yes | caudatus_Native |
| PI 490609 | PI490609 | caudatus | ECU | Native | 4.3X | 47.5% | 17.2 | Yes | caudatus_Native |
| PI 490612 | PI490612 | caudatus | PER | Native | 12.8X | 97.5% | 10 | Yes | caudatus_Native |
| PI 490673 | PI490673 | quitensis | ECU | Native | 11.1X | 97.0% | 10.1 | Yes | quitensis_Native |
| PI 490705 | PI490705 | quitensis | ECU | Native | 11.9X | 97.5% | 10.1 | Yes | quitensis_Native |
| PI 490720 | PI490720 | quitensis | ECU | Native | 14.5X | 97.3% | 9.6 | Yes | quitensis_Native |
| PI 511679 | PI511679 | caudatus | ARG | Native | 9.0X | 96.7% | 10.8 | Yes | caudatus_Native |
| PI 511680 | PI511680 | caudatus | ARG | Native | 9.2X | 97.2% | 10.5 | Yes | caudatus_Native |
| PI 511681 | PI511681 | caudatus | BOL | Native | 9.1X | 97.7% | 10.4 | Yes | caudatus_Native |
| PI 511686 | PI511686 | caudatus | PER | Native | 10.9X | 97.1% | 10.4 | Yes | caudatus_Native |
| PI 511687 | PI511687 | caudatus | PER | Native | 12.3X | 96.9% | 10.1 | Yes | caudatus_Native |
| PI 511690 | PI511690 | caudatus | PER | Native | 9.4X | 96.6% | 11.1 | Yes | caudatus_Native |
| PI 511696 | PI511696 | caudatus | PER | Native | 10.1X | 95.8% | 10.4 | Yes | caudatus_Native |
| PI 511704 | PI511704 | caudatus | PER | Native | 12.2X | 97.2% | 9.8 | Yes | caudatus_Native |
| PI 511706 | PI511706 | caudatus | PER | Native | 9.6X | 96.2% | 11 | Yes | caudatus_Native |
| PI 511712 | PI511712 | caudatus | ECU | Native | 9.9X | 96.9% | 10.6 | Yes | caudatus_Native |
| PI 511713 | PI511713 | cruentus | PER | Native | 10.9X | 95.9% | 9.6 | Yes | cruentus_Native |
| PI 511714 | PI511714 | cruentus | PER | Native | 8.4X | 97.4% | 9.1 | Yes | cruentus_Native |
| PI 511717 | PI511717 | cruentus | GTM | Native | 9.7X | 97.0% | 10.4 | Yes | cruentus_Native |
| PI 511724 | PI511724 | hybridus | MEX | Native | 9.7X | 95.8% | 10.6 | Yes | hybridus_CA_Native |
| PI 511731 | PI511731 | hypochondriacus | MEX | Native | 3.1X | 24.6% | 18.5 | Yes | hypochondriacus_Native |
| PI 511736 | PI511736 | quitensis | BOL | Native | 9.7X | 96.4% | 9.4 | Yes | quitensis_Native |
| PI 511737 | PI511737 | quitensis | ECU | Native | 12.4X | 97.3% | 10 | Yes | quitensis_Native |
| PI 511741 | PI511741 | quitensis | ECU | Native | 13.0X | 91.5% | 9.6 | Yes | quitensis_Native |
| PI 511745 | PI511745 | quitensis | ECU | Native | 10.2X | 97.1% | 10.7 | Yes | quitensis_Native |
| PI 511747 | PI511747 | quitensis | ECU | Native | 9.7X | 97.6% | 10.4 | Yes | quitensis_Native |
| PI 511749 | PI511749 | quitensis | ECU | Native | 11.5X | 97.3% | 10.1 | Yes | quitensis_Native |

| | | | | | | | | | |
|---|---|---|---|---|---|---|---|---|---|
| PI 511754 | PI511754 | hybridus | ECU | Native | 13.5X | 94.6% | 8.8 | Yes | hybridus_SA_Native |
| PI 576481 | PI576481 | cruentus | MEX | Native | 10.2X | 97.3% | 9.2 | Yes | cruentus_Native |
| PI 576482 | PI576482 | cruentus | MEX | Native | 9.9X | 97.3% | 10.1 | Yes | cruentus_Native |
| PI 604568 | PI604568 | hybridus | MEX | Native | 8.3X | 96.0% | 16.5 | Yes | hybridus_SA_Native |
| PI 604574 | PI604574 | hybridus | MEX | Native | 11.6X | 97.1% | 9.5 | Yes | hybridus_CA_Native |
| PI 604581 | PI604581 | hypochondriacus | MEX | Native | 11.5X | 98.6% | 4.5 | Yes | hypochondriacus_Native |
| PI 604582 | PI604582 | hybridus | MEX | Native | 2.4X | 23.5% | 30.6 | Yes | hybridus_CA_Native |
| PI 604587 | PI604587 | hypochondriacus | MEX | Native | 11.3X | 97.1% | 5 | Yes | hypochondriacus_Native |
| PI 604595 | PI604595 | hypochondriacus | MEX | Native | 9.8X | 97.6% | 3.6 | Yes | hypochondriacus_Native |
| PI 606798 | PI606798 | cruentus | MEX | Native | 9.5X | 97.0% | 10.1 | Yes | cruentus_Native |
| PI 608019 | PI608019 | caudatus | ECU | Native | 10.5X | 96.0% | 10.5 | Yes | caudatus_Native |
| PI 633589 | PI633589 | hypochondriacus | MEX | Native | 11.1X | 98.1% | 4.3 | Yes | hypochondriacus_Native |
| PI 636180 | PI636180 | hybridus | COL | Native | 6.6X | 89.6% | 17.3 | Yes | hybridus_SA_Native |
| PI 642741 | PI642741 | caudatus | BOL | Native | 0.9X | 94.9% | 11.6 | Yes | caudatus_Native |
| PI 643036 | PI643036 | hypochondriacus | MEX | Native | 12.0X | 98.0% | 3.2 | Yes | hypochondriacus_Native |
| PI 643037 | PI643037 | cruentus | MEX | Native | 8.6X | 97.7% | 9.8 | Yes | cruentus_Native |
| PI 643039 | PI643039 | cruentus | MEX | Native | 12.0X | 96.6% | 9.3 | Yes | cruentus_Native |
| PI 643041 | PI643041 | hypochondriacus | MEX | Native | 12.3X | 98.0% | 3.5 | Yes | hypochondriacus_Native |
| PI 643049 | PI643049 | cruentus | MEX | Native | 11.8X | 96.5% | 9.5 | Yes | cruentus_Native |
| PI 643058 | PI643058 | cruentus | MEX | Native | 13.2X | 97.1% | 8.9 | Yes | cruentus_Native |
| PI 643067 | PI643067 | hypochondriacus | MEX | Native | 11.4X | 97.0% | 3.7 | Yes | hypochondriacus_Native |
| PI 643070 | PI643070 | hypochondriacus | MEX | Native | 13.2X | 98.2% | 2.8 | Yes | hypochondriacus_Native |
| PI 649217 | PI649217 | caudatus | PER | Native | 9.3X | 96.5% | 11 | Yes | caudatus_Native |
| PI 649227 | PI649227 | caudatus | PER | Native | 13.1X | 95.0% | 9.3 | Yes | caudatus_Native |
| PI 649228 | PI649228 | caudatus | PER | Native | 8.9X | 95.8% | 11.1 | Yes | caudatus_Native |
| PI 649230 | PI649230 | caudatus | PER | Native | 9.3X | 97.0% | 10.7 | Yes | caudatus_Native |
| PI 649509 | PI649509 | cruentus | MEX | Native | 9.7X | 97.3% | 10.1 | Yes | cruentus_Native |
| PI 649514 | PI649514 | cruentus | MEX | Native | 7.0X | 62.8% | 11.6 | Yes | cruentus_Native |
| PI 649524 | PI649524 | cruentus | MEX | Native | 10.4X | 96.4% | 9.7 | Yes | cruentus_Native |
| PI 649529 | PI649529 | hypochondriacus | MEX | Native | 9.4X | 97.3% | 3.2 | Yes | hypochondriacus_Native |
| PI 649537 | PI649537 | hypochondriacus | MEX | Native | 12.0X | 97.9% | 3.6 | Yes | hypochondriacus_Native |
| PI 649559 | PI649559 | hypochondriacus | MEX | Native | 10.8X | 97.1% | 3.8 | Yes | hypochondriacus_Native |
| PI 649575 | PI649575 | hypochondriacus | MEX | Native | 10.5X | 97.9% | 3.5 | Yes | hypochondriacus_Native |
| PI 649602 | PI649602 | hypochondriacus | MEX | Native | 11.8X | 97.8% | 2.5 | Yes | hypochondriacus_Native |
| PI 649607 | PI649607 | hypochondriacus | MEX | Native | 11.5X | 97.9% | 3.7 | Yes | hypochondriacus_Native |
| PI 649609 | PI649609 | cruentus | MEX | Native | 9.6X | 93.5% | 10 | Yes | cruentus_Native |
| PI 649623 | PI649623 | hypochondriacus | MEX | Native | 10.9X | 97.4% | 2.9 | Yes | hypochondriacus_Native |
| PI 658728 | PI658728 | cruentus | MEX | Native | 9.5X | 97.0% | 9.8 | Yes | cruentus_Native |
| PI 667158 | PI667158 | hybridus | GTM | Native | 10.1X | 97.0% | 7 | Yes | hybridus_CA_Native |
| PI 667160 | PI667160 | cruentus | GTM | Native | 9.0X | 96.4% | 10.3 | Yes | cruentus_Native |
| PI 667165 | PI667165 | cruentus | BRA | Native | 9.7X | 97.0% | 10.2 | Yes | cruentus_Native |

**Table S2** List of genes annotated in 10kb up-downstream region of assocaited GWAS SNP.

| Chromosome | Start | End | Name | Description |
|---|---|---|---|---|
| 1 | 54290 | 74290 | AHp000004 | lysine-specific demethylase |
| 4 | 2539899 | 2559899 | AHp006724 | L-galactono-1,4-lactone dehydrogenase |
| 5 | 18969424 | 18989424 | AHp009270 | malic enzyme |
| 9 | 20373084 | 20393084 | AHp015126 | isoform X1 |
| 10 | 18379508 | 18399508 | AHp016221 | Signal peptidase complex subunit |
| 10 | 18379508 | 18399508 | AHp016222 | DOMON domain-containing protein |
| 11 | 2043844 | 2063844 | AHp016706 | Belongs to the protein kinase superfamily. Ser Thr protein kinase family |
| 11 | 22024217 | 22044217 | AHp017947 | Boron transporter |
| 11 | 22024217 | 22044217 | AHp017948 | Catalyzes the phosphorylation of pantothenate the first step in CoA biosynthesis. May play a role in the physiological regulation of the intracellular CoA concentration |
| 12 | 16842318 | 16862318 | AHp019109 | Domain of unknown function (DUF4336) |
| 12 | 16842318 | 16862318 | AHp019110 | ribulose-phosphate 3-epimerase |
| 13 | 5600536 | 5620536 | AHp019501 | disulfide-isomerase |
| 15 | 9943938 | 9963938 | AHp022473 | histone H2B |
| 15 | 9943938 | 9963938 | AHp022474 | histone H2B |

**Table S3** 95% confidence interval along mean nucleotide diversity estimated 50 times by sub-sampling 10 individuals from each population at a time using angsd.

| Species | Region | 95% CI along mean Pi | |
|---|---|---|---|
| | | LowerBound | UpperBound |
| *A. hybridus* | Wild | 0.008589 | 0.008593 |
| *A. quitensis* | | 0.001854 | 0.00414 |
| *A. caudatus* | Native | 0.002349 | 0.002955 |
| *A. cruentus* | | 0.002241 | 0.005253 |
| *A. hypochondriacus* | | 0.002584 | 0.005005 |
| *A. caudatus* | India | 0.003213 | 0.00473 |
| *A. cruentus* | | 0.003984 | 0.003985 |
| *A. hypochondriacus* | | 0.003243 | 0.004436 |
| Hypochondriacus-mix | | 0.002665 | 0.003784 |
| Hybrid | | 0.003125 | 0.003513 |

**Table S4** Parameter estimates of best-fit demographic model for the two species with 95% confidence interval.

| **Native Bottleneck- introduced expansion- continuous gene flow** | | | |
|---|---|---|---|
| *A. hypochondriacus* | | | |
| **Parameters** | **PointEstimates** | **LowerBound-05** | **UpperBound-95** |
| **ANCSIZE** | 14693 | 10470 | 130783 |
| **NATIVE** | 1615042 | 10153 | 8543408 |
| **NPOPOUT** | 277553 | 120469 | 973019 |
| **NBOT** | 1355 | 1015 | 764414 |
| **INDIA** | 38513 | 27919 | 341274 |
| **TDIV** | 1240 | 908 | 9844 |
| **TBOT** | 15 | 11 | 8627 |
| **R1** | -0.00562649 | -0.007668 | -0.000707 |
| **MIG12** | 0.0011074 | 0.000142 | 0.001467 |
| **MIG21** | 0.000189159 | 2.3E-05 | 0.000265 |
| **TDUR** | 653 | 132 | 905 |
| **TENDBOT** | 667 | 382 | 9008 |
| **MaxEstLhood** | -6009456.677 | -108604.808 | -105911.954 |
| **MaxObsLhood** | -5936545.021 | -108356.491 | -105788.505 |

| **Native Bottleneck- introduced expansion- Introgression with unknown population** | | | |
|---|---|---|---|
| *A. caudatus* | | | |
| **Parameters** | **Point Estimates** | **LowerBound-05** | **UpperBound-95** |
| **ANCSIZE** | 649899 | 384442 | 1040340 |
| **NATIVE** | 3069064 | 32756 | 9509859 |
| **NPOPOUT** | 243883 | 111614 | 910672 |
| **NBOT** | 1108 | 1012 | 1979 |
| **INDIA** | 9868060 | 5645064 | 12346802 |
| **TDIV** | 2666 | 1260 | 6051 |
| **TBOT** | 383 | 137 | 648 |
| **R1** | -0.00416919 | -0.009469 | -0.001713 |
| **MIG13** | 0.0020243 | 1.9E-05 | 0.003223 |
| **TDUR** | 181 | 158 | 320 |
| **TENDBOT** | 565 | 333 | 934 |
| **MaxEstLhood** | -5030246.35 | -987420.829 | -980605.148 |
| **MaxObsLhood** | -4970605.226 | -987286.588 | -980480.136 |

**Table S5** List of genes in the putative sweep regions identified using XP-CLR.

**Table S6** Stepwise forward variable selection procedure using ordiR2step. The stopping criteria used were - variable significance of $p < 0.01$ using 1000 permutations, and the adjusted $R^2$ of the global model.

| **Variable** | **R square** | **Cumulative R square** | **F value** | **P value** |
|---|---|---|---|---|
| *A. caudatus* | | | | |
| Bio3 | 0.108 | 0.108 | 6.1092 | 0.002 ** |
| Bio9 | 0.009 | 0.117 | 1.423 | 0.018 * |
| *A. hypochondriacus* | | | | |
| Bio2 | 0.094 | 0.094 | 4.3075 | 0.002 ** |
| Bio7 | 0.014 | 0.109 | 1.5251 | 0.004 ** |

**Table S7** Full and partial Redundancy analysis (RDA) to partition variances into genetic, climatic and geographic components.

| Partial RDA models | Inertia | R 2 | p (>F) | Proportion of explainable variance | Proportion of total variance |
|---|---|---|---|---|---|
| ***A. caudatus*** | | | | | |
| Full model: F ∼clim. + geog. + struct. | 4076.4 | 0.38 | 0.001 *** | 1 | 0.31 |
| Pure climate: F ∼clim. ∣ (geog. + struct.) | 423.8 | 0.039 | 0.24 | 0.1 | 0.032 |
| Pure structure: F ∼struct. ∣ (clim. + geog.) | 1423.7 | 0.13 | 0.001 *** | 0.34 | 0.108 |
| Pure geography: F ∼geog. ∣ (clim. + struct.) | 617.2 | 0.057 | 0.244 | 0.15 | 0.046 |
| Total unexplained | 6635.4 | | | | 0.5 |
| Total inertia | 13176.5 | | | | 1 |
| ***A. hypochondriacus*** | | | | | |
| Full model: F ∼clim. + geog. + struct. | 1252.6 | 0.413 | 0.001 *** | 1 | 0.32 |
| Pure climate: F ∼clim. ∣ (geog. + struct.) | 170.35 | 0.056 | 0.118 | 0.136 | 0.044 |
| Pure structure: F ∼struct. ∣ (clim. + geog.) | 474.35 | 0.157 | 0.001 *** | 0.379 | 0.121 |
| Pure geography: F ∼geog. ∣ (clim. + struct.) | 237.78 | 0.079 | 0.223 | 0.19 | 0.061 |
| Total unexplained | 1773.4 | | | | 0.454 |
| Total inertia | 3908.48 | | | | 1 |

**Table S8** List of annotated genes in the 20kb up-downstream regions of significantly associated loci identified using RDA
